# A chromosome-scale genome assembly of *Hordeum erectifolium*: genomic, transcriptomic and anatomical adaptations to drought in a wild barley relative

**DOI:** 10.1101/2025.08.27.672388

**Authors:** Einar Baldvin Haraldsson, Michael Anokye, Thea Rütjes, Helena Toegelová, Zuzana Tulpová, Hana Šimková, Jia-Wu Feng, Martin Mascher, Maria von Korff

## Abstract

- Wild crop relatives are valuable genetic resources for improving stress adaptation in cultivated species, but their effective use depends on high-quality reference genomes integrated with phenotypic and molecular datasets. *Hordeum erectifolium*, a wild relative of barley (*H. vulgare*), is adapted to intermittent and prolonged drought and saline soils, making it an excellent model for stress-adaptation research.
- We assembled a chromosome-scale, annotated reference genome of *H. erectifolium* comprising 3.85 Gbp, and identified 71,475 genes supported by a tissue-specific gene expression atlas. Comparative morphological, physiological, and transcriptomic analyses under water limitation were conducted with cultivated and wild barley.
- *H. erectifolium* displayed a greater density of leaf veins and sclerenchyma cells, alongside rapid leaf rolling upon dehydration. Genomic comparisons revealed structural variations, independent transposon-driven evolution, and copy number expansions of desiccation-responsive gene families relative to barley. The transcriptional responses of *H. erectifolium* and barley to water limitation suggested contrasting drought-adaptation strategies: metabolic down-regulation and survival prioritization in *H. erectifolium* versus maintenance of metabolic activity and competitiveness in barley.
- Our data suggest that *H. erectifolium* is genetically primed for survival under drought through anatomical adaptations, gene family expansion, efficient shutdown of growth-related metabolism, and rapid recovery upon rehydration.

## Introduction

To ensure food security in the face of a changing climate, it is essential to develop and implement innovative strategies that enhance crop resilience to extreme weather events, diversify food sources, and optimize resource use. Crop wild relatives (CWR), the close wild relatives of domesticated crop species, harbour novel traits and alleles that serve as a valuable resource for breeding stress-resilient crops and for advancing research into the genetic and genomic basis of plant adaptation (Feuillet et al., 2008; Bohra et al., 2022).

CWRs are categorized into three gene pools. The primary gene pool includes the cultivated crop and its closest wild relatives that can cross easily and produce fertile offspring. The secondary relatives can be crossed with some difficulty and reduced fertility, while tertiary relatives require advanced methods like genetic engineering or wide hybridization to overcome strong reproductive barriers (Harlan and de Wet, 1971; Baker et al., 2020; Kashyap et al., 2022). For over a century, CWR have been used for crop improvements, in particular as a resource for disease and pest resistance genes, but also to improve resistance to abiotic stresses (Hajjar and Hodgkin, 2007; Renzi et al., 2022). The introgression of stress escape and avoidance strategies into elite germplasm has often been achieved by introducing exotic alleles altering phenology, the timing of germination and reproductive development (Duc et al. 2015; Farooq et al., 2025 ). More recently, CWRs have been used to introduce perennial growth, a complex trait that extends survival and allows for reproduction over many seasons, into annual crops (Gruner and Miedaner, 2021; Zhang et al., 2023). Perennial crops promise to reduce labor costs, decrease input requirements, reduce soil erosion and enhance soil health, thereby supporting more sustainable agriculture (Zhang et al., 2023). However, the introgression of novel traits such as perenniality, disease and abiotic stress resistance from CWR is hampered by hybridization barriers, low fertility, and time and labor costs for generating advanced lines (Wendler et al., 2015). The availability of reference genomes and genetic tools for gene identification and transfer now promises to deliver novel, efficient approaches for using CWR in research and breeding (Brozynska et al., 2016; Gao et al., 2024).

The annual crop barley (*Hordeum vulgare* ssp. *vulgare*) was domesticated from the wild barley ancestor *H. vulgare ssp. spontaneum* in the Fertile Crescent (Badr et al., 2000, Pankin et al. 2018). Wild and cultivated barley belong to the primary gene pool and are characterised by natural continuous gene flow in the Fertile Crescent, where wild and cultivated barley co-occur (Pankin et al. 2018). Wild barley has been successfully used to introgress novel alleles for disease resistance, abiotic stress tolerance and flowering time into elite barley (Matus et al. 2003; von Korff et al. 2004; Hernandez et al. 2020). In addition, *H. bulbosum* from the secondary CWR gene pool was a source of resistance genes to powdery mildew, leaf rust, barley mild and yellow mosaic virus for breeding elite barley (Walther et al., 2000, Vincent et al., 2012). However, CWRs from the tertiary gene pool within the *Hordeum* genus have not yet been used for barley improvement, not only because of hybridization barriers but also because of a lack of genetic and genomic resources. The *Hordeum* genus, comprising about 33 annual and perennial species, originated in the Mediterranean some 12 million years ago (Mya) and has since migrated globally, with 16 species in South America, which have evolved in the last 1.5 Mya (Blattner, 2018). The South American clade of the *Hordeum* genus contains closely related species adapted to different ecological niches and with different life history strategies, making them an interesting resource for trait diversification and research into the physiological and genetic underpinnings of adaptation (Bothmer and Komatsuda, 2010). Among the South American clade, *Hordeum erectifolium* is one of the younger species, with the last common ancestor of *H. erectifolium* and its closest sister species, *H. stenostachys,* dating back to ca. 1.3 Mya, whereas the split of the branches separating *H. erectifolium* and *H. vulgare* occurred approximately 9.2 Mya (Brassac & Blattner, 2015). *H. erectifolium* is endemic to the southernmost part of the semiarid Pampas, Argentina, where it grows among halophytes near a salt lake (Bothmer et al., 1985). This region is impacted by frequent irregular periods of drought that can extend to up to a year (Bothmer et al., 1985; Cai et al., 2020; Sgroi et al., 2021; WMO, 2020). Due to its small distribution area, it is considered a near-threatened species (IUCN 2024) and is listed as a high priority for *ex-situ* conservation within the *Hordeum* genus; only a single accession of *H. erectifolium* is available in gene banks (Vincent et al., 2012). Being a perennial species, *H. erectifolium* has to persist through long and unpredictable periods of drought and saline soil. It has thus evolved unique stress avoidance traits, such as very glaucous erect basal leaves and pubescence on the leaf abaxial side, a thick wax coating, and large suberized silica cells (Bothmer et al., 1985, 1995). These unique adaptations make *H. erectifolium* an interesting genetic resource to explore the physiological and genetic underpinnings of stress adaptation compared to the closely related cultivated crop barley.

We present a chromosome-scale, annotated reference genome of *H. erectifolium*, together with a comprehensive tissue-specific gene expression atlas. By comparing genomic, transcriptomic, and morphological responses to water limitation with those of cultivated barley, we highlight stress-adaptive strategies employed by *H. erectifolium*. Our findings underscore the potential of this perennial close relative of barley as a model for dissecting the morphological, physiological, and genetic bases of stress adaptation.

## Materials and methods

Here is a brief description of methods and material, a more details are found in the Online methods and materials (Online methods).

### Plant material and growth conditions

For all experiments we used single descent propagated seeds of *H. erectifolium* acc. NGB6816 (Nordic Genetic Resource Center, Sweden), and *H. v. spontaneum* acc. B1K-04-12 from the Barley1K collection (Hübner et al., 2009), *H. v. spontaneum* acc. HID-4 (Liller et al., 2017), and *H. vulgare* cv. Morex. *H. erectifolium* and Morex were used in all experiments, but B1K-01-12 was used for leaf anatomical phenotyping, and HID-4 for specific leaf area (SLA) and elemental carbon and nitrogen measurements. Unless specified otherwise, plants were consistently grown under the following conditions. Seeds were sown in a mixture of 93 % (v/v) Einheitserde ED73 (Einheitserdewerke Werkverband e.V., Sinntal-Altengronau, Germany), 6.6 % (v/v) sand, and 0.4 % (v/v) Osmocote exact standard 3-4M (Scotts Company LLC). Stratified at 4 °C before being placed in a growth chamber with long day conditions (16 h light, 8 h dark, at 20 °C day/16 °C night, 60 % relative humidity) for germination. Ten days after germination, they were then vernalized for 8 weeks at 4 °C under short day conditions (8 h light, 16 h dark, at 4 °C day/4 °C night), before being transferred back to the long day conditions. Plants used for specific leaf area and elemental carbon and nitrogen measurements were cultivated as described above. After vernalization, they were repotted to 7.5 L pots and grown further in a common garden at the Botanical Garden of the Heinrich-Heine-Universität Düsseldorf (HHU), data collected were from three summer seasons of 2021 – 2023.

### Phenotyping

To visualize the leaf anatomy, we used toluidine blue to stain the lignin of transversely cut leaves. We collected the flag leaf and the leaf below the flag leaf, and 1 cm sections at the midpoint of the leaf were thinly sliced and fixed for staining with toluidine blue for microscope image capture. The specific leaf area (SLA) was measured by scanning the area (cm^2^) of a main culm flag leaf with Petiole (Petiole LTD, U.S.A.), an application on a mobile device. The dry weight (mg) was measured after drying at 65 °C, and SLA was calculated as SLA = cm^2^ / mg. We measured elemental carbon and nitrogen by pooling the flag leaves from the first three reproductive tillers per plant at the grain filling stage. They were dried at 65 °C before being pulverized to a fine powder, and approximately 2 mg were accurately measured. The samples were provided to the Metabolomics and Metabolism Laboratory (CMML, Heinrich-Heine University Düsseldorf, Germany) for measurements of elemental carbon and nitrogen.

### Generation of an annotated chromosome-scale genome assembly

A single plant of *H. erectifolium* acc. NGB6816 grown under control conditions, was used for extracting high molecular weight (HMW) DNA. HMW DNA was extracted from young leaf material and sequenced on 27 Oxford Nanopore Technology (ONT) flow cells (Table **S1**), and with 10x Genomics Linked-Reads sequencing for assembly polishing and the first scaffolding. Bionano optical genome mapping was made with material of the same plant that also provided the seeds used to generate seedlings for Hi-C chromosome conformation capture sequencing. We processed the 10x Genomics Linked-Reads data with Long Ranger (10x Genomics, U.S.A.), generating paired-end Illumina (PE150) short-reads data with Linked-Reads information in headers (short-reads). We performed initial genome kmer-based characterization (size, ploidy level, heterozygosity), by analyzing the short-reads data using Jellyfish and findGSE (Marçais & Kingsford, 2011; Sun et al., 2018).

The ONT sequencing was basecalled with Guppy (ONT, United Kingdom) using a quality threshold of ≥Q7 and subsequently trimmed sequencing adapters with Porechop (Wick et al., 2017), before assembling the ONT long-reads with Flye (Kolmogorov et al., 2019). The assembly was progressively polished; first, we used minimap2 to map the ONT long-reads to the assembly before each polishing step, first with Racon and then with Medaka (ONT, United Kingdom) (Vaser et al., 2017; Li, 2018). Thereafter, we mapped the short-reads with BWA-MEM2 to the long-read polished assembly and performed two rounds of polishing with Hapo-G and repeated the mapping between rounds (Vasimuddin et al., 2019; Aury & Istace, 2021). The polished assembly was progressively scaffolded from contigs to pseudomolecules. Initially, Tigmint was used with the 10x Genomics Linked-Reads short-read data (Jackman et al., 2018), followed by hybrid-scaffolding using Bionano optical genome mapping (Šimková et al., 2023). Finally, the assembly was organized into pseudomolecules using the TRITEX pipeline with Hi-C chromosome conformation capture sequencing (Monat et al., 2019). Assembly quality, completeness, and metrics were assessed at each step using complementary tools: Merqury for k-mer–based evaluation of assembly accuracy and k-mer completeness, BUSCO for gene space completeness, and QUAST for standard assembly statistics (Mikheenko et al., 2018; Rhie et al., 2020; Manni et al., 2021). A unique genome assembly identifier, lpHorErec1.1, was registered at ToLID – Tree of Life Identifiers, id.tol.sanger.ac.uk.

We conducted a comprehensive tissue-time specific transcriptome profiling of *H. erectifolium*. All mature RNA plant tissue samples and seeds used for seedlings and germinating seed samples were collected from the same plant as was previously harvested for HMW DNA. We selected twelve tissues, ten of which were sampled at two time points during the day, in the morning (MOR, ZT 1-3) and evening (EVE, ZT 13-16), a total of 22 samples. The vegetative tissues contained three-day-old germinating seeds (GS3), whole roots (RO8, 8 days post germination), and whole shoots (LS8, 8 days post germination). At anthesis, the third nodes (NOD), fourth internodes (INT), and flag leaves (FLF) were collected. The six reproductive samples consisted of developing spikes at the spikelet induction phase (ESP, W3.0-4.5), at floral development (MSP, W5.0-6.5), and during rapid spike and floret growth (LSP, W7.0-8.0), anthers (ANT) and ovules (OVU) were individually dissected at flowering (W10) and caryopses (CAR, 10 days post anthesis) (Waddington et al., 1983). RNA was extracted, and each tissue was barcoded and sequenced with PacBio IsoSeq (Pacific Biosciences, U.S.A).

For the gene annotation, we first mapped the 22 PacBio IsoSeq samples to the assembled genome and processed them with the isoseq3 pipeline (Pacific Biosciences, U.S.A). We predicted the open reading frames (ORF) for each isoform using TransDecoder (Haas, 2023), and additional *ab initio* structural gene annotation was made with Helixer (Holst et al., 2023). The IsoSeq-Transdecoder and Helixer results were merged for a final gene annotation, and functionally annotated them with InterProScan and Mercator4 (Jones et al., 2014; Bolger et al., 2021). We further predicted the presence of lncRNA in the IsoSeq data. Transcript abundance in 22 tissue- and time-specific samples was quantified as normalized transcripts per million (TPM) using IsoQuant (Prjibelskwe et al., 2023), and the samples were clustered using Principal Component Analysis (PCA). We visualized the top 20 contributing genes, separating the 22 tissues along the principal components 1 (PC1) and PC2.

For genomic and genetic comparisons between *H. erectifolium* acc. NGB6816 and barley, we used the barley reference genome MorexV3, and the genome of a wild barley *H. v. spontaneum* acc. B1K-04-12 (Jayakodi et al., 2020; Mascher et al., 2021). We annotated transposable elements (TE) and repetitive regions of the *H. erectifolium* and barley genomes with EDTA and further classified LTR superfamilies with TEsorter (Ou et al., 2019; Zhang et al., 2022). The telomeric ends were identified by aligning the telomeric sequence TTTAGGG^8^ and centromeres using the *CRM* coding domains, extracted with TEsorter from the EDTA annotation of LTR retrotransposon (Ou et al., 2019; Zhang et al., 2022). The sequences were aligned with BLASTN to the three genomes (Camacho et al., 2009). We identified the centromere midpoints from characteristic alignment density patterns using hdrcde (Hyndman et al., 2023). We calculated the chromosomal synteny among the genomes by mapping them against one another with minimap2 (Li, 2018), and structural variations were calculated with Synteny and Rearrangement Identifier (SyRI) before visualization with plotsr (Goel et al., 2019; Goel & Schneeberger, 2022).

### Gene family evolution analysis with Orthofinder and CAFE5

A study of unique and shared genes and gene families was made between *H. erectifolium* and eight additional species, including the two barley genotypes. The proteomes of six species were retrieved from the JGI Phytozome database: *Arabidopsis thaliana, Sorghum bicolor, Zea mays, Oryza sativa, Brachypodium distachyon,* and *Triticum aestivum* cv. Chinese Spring (Goodstein et al., 2012). We assigned the proteomes to Hierarchical Orthologous Groups (HOG) with Orthofinder, and gene family evolution analyzed with CAFE5 (Emms & Kelly, 2019; Mendes et al., 2021), with a species separation time of 59 million years retrieved from TimeTree5 ( Kumar et al., 2022).

### Cross-species drydown experimental setup and differential gene expression analysis

We set up a drydown and recovery experiment to study the cross-species transcriptomic and physiological responses to a reduction in relative leaf water content. One seed was sown per 7×7×8 cm pot with 150±1 g of soil, and plants were germinated and further cultivated under 12 h light/ 12 h dark and temperatures 20 °C day/16 °C night, 60 % relative humidity. *H. erectifolium* plants were grown for eight weeks, and Morex plants for two weeks to ensure comparable biomass at the start of the treatment. Soil moisture was adjusted to 50 % field capacity (FC), control FC, and water was withheld for six days before rewatering to control FC. Four replicate samples of the second leaves from the main tillers were collected for transcriptomic analyses at four days of treatment (DOT) at ZT 8, on DOT 2, 5, 6, and 7, 24 hours after re-watering. Leaf relative water content (RWC) was measured at each DOT. Total RNA was extracted, and the transcriptome was Illumina PE150 sequenced.

The RNAseq reads were mapped to their respective genome with STAR, and quantification was done using featureCounts (Dobin et al., 2013; Liao et al., 2014). The quantified reads were processed and normalized with edgeR, and differentially expressed genes (DEG) analyzed at individual timepoints with edgeR. maSigPro was used for time-course and co-expression analysis of DEGs in response to treatment over time (Nueda et al., 2014; Chen et al., 2024). Gene Ontology (GO) functional enrichment was made with Clusterprofiler (Xu et al., 2024). Single copy orthologs (SCO) between *H. erectifolium* and Morex were retrieved from HOGs and were used for direct DEG comparison between species.

### Data analysis

Data wrangling, statistics, and visualization was done in R (v. 4.4.3) (R Core Team, 2024) were made with tidyverse (v 2.0.0), and statistics with agricole (v. 1.5) in (Wickham et al., 2019; Mendiburu, 2023).

## Results

### Leaf morphology of *H. erectifolium*, cultivated and wild barley

*H. erectifolium* leaves were erect and had a glaucous appearance (Fig. **1a,b**). We observed that upon reduced water availability, *H. erectifolium* rolled its leaves, whereas wild and cultivated barley maintained flat leaves (Fig. **1b,c**). Leaf rolling is typically supported by specific leaf anatomies, such as reduced or a lack of bulliform cells that provide structural support (Redmann, 1985). We therefore compared the leaf anatomy of the flag leaf and the first leaf below the flag leaf of *H. erectifolium*, cultivated (Morex), and wild barley. Here we present the results for the flag leaf, which were confirmed in the first leaf below (Fig. **S1**). The flag leaf was characterised by a prominent ribbed leaf structure with short and long trichomes on the adaxial side, with no discernible bulliform cells. We also observed that as the flag leaf of *H. erectifolium* develops and extends, the abaxial side generally faces upwards. The white appearance of the flag leaf indicated a layer of cuticular wax (Fig. **1a,b**). In contrast, cultivated and wild barley both had a flat leaf surface, much shorter trichomes, and pronounced bulliform cells (Fig. **1d**). We quantified the traverse vein density, the minor: major vein ratio, leaf thickness and width, as well as specific leaf area and elemental carbon: nitrogen ratio. Major veins are distinguished from minor veins by their additional bundle sheath extensions (BSE) (Perico et al., 2022). The average number of veins in *H. erectifolium* per mm was 6.2 vs 3.4 and 3.9 veins/mm, in *H. erectifolium* versus cultivated and wild barley genotypes, respectively (Fig. **1e**, **S1**). The minor: major vein ratio had significantly shifted in *H. erectifolium* compared to cultivated and wild barley, from 1.3 vs. 3.0 and 3.1, respectively. While cultivated and wild barley had three minor veins flanked by major veins, the central minor vein had become a major vein in *H. erectifolium* (Fig. **1d,f**, **S1**). We noted that *H. erectifolium* had overall more extensive sclerenchyma cells within their BSE on the adaxial side of the veins compared to barley. Sclerenchyma cells are connected with increased photosynthetic activity, water transport, and more rapid stomatal closure in addition to structural support (Buckley et al., 2011). We found that Specific Leaf Area (SLA) was significantly lower and carbon: nitrogen (C:N) ratios were significantly higher in *H. erectifolium* than in cultivated and wild barley; SLA with 1.5 mm^2^/mg vs 2.4 mm/mg and 2.2 mm/mg, and C:N ratio of 13 vs 7.0 and 7.8, in *H. erectifolium and* cultivated and wild barley, respectively (Fig. **1g,h**). Low SLA and high C:N ratio indicate a slow investment in leaf growth and the photosynthetic apparatus, and thus a conservative growth strategy with improved resource use efficiency (Poorter et al., 2009; Onoda et al., 2017). Leaf thickness and width were overall not different between species; however, the leaf width of Morex was significantly greater than that of wild barley and *H. erectifolium* (Fig. **1i,j**, **S1**). In short, *H. erectifolium* exhibited significantly higher leaf venation and a greater number of major veins—likely due to the conversion of minor veins into major ones—as well as a higher abundance of sclerenchyma cells on both the abaxial and adaxial sides of the vascular bundles, reflecting greater photosynthetic tissue compartmentalization and water distribution efficiency.

**Fig. 1:**
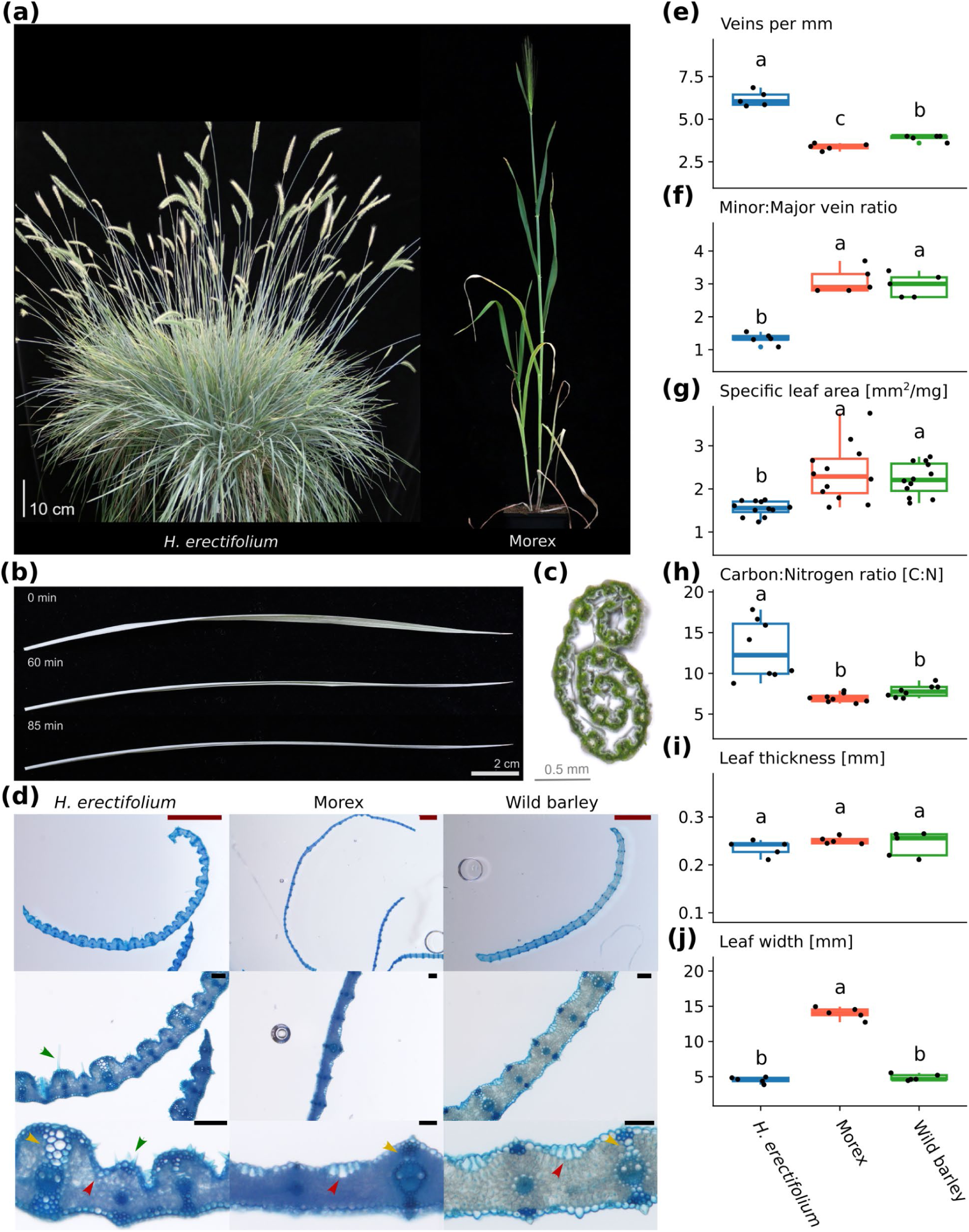
Leaf morphology of *H. erectifolium*, cultivated (Morex) and wild barley. **(a)** A representative of *Hordeum erectifolium* (left), accession NGB 6816, from which all genetic and transcriptomic data was collected, and an example of *H. vulgare cv.* Morex (right). **(b)** Abscised leaf of *H. erectifolium*, folding inward towards the adaxial side of the midriff. **(c)** Transverse cut of a *H. erectifolium* leaf, rolled towards the adaxial side of the midriff. **(d)** Transverse cuts of *H. erectifolium*, Morex, and wild barley, lignin of fixated tissue was stained with toluidine blue. Red arrows point at bulliform cells in Morex and wild barley, and the expected location in *H. erectifolium*. Trichomes are indicated by green arrows in *H. erectifolium*. Yellow arrows point at the adaxial bundle sheath extensions (BSE) on a major vein; the BSE is as well on the abaxial side of the vein. Scale bars for 1000 µm are indicated in red and 100 µm in black. Quantitative leaf characteristics of the flag leaf (FL) in *H. erectifolium,* Morex, and wild barley: **(e)** number of veins per mm, n = 5, **(f)** minor to major vein ration, n = 5, **(g)** specific leaf area, cm^2^/mg, n = 8, **(h)** ratio of elemental carbon and nitrogen, n = 8. **(i, j)** FL thickness and width, mm, n = 5. Different letters indicate significantly differing groups, ANOVA with post-hoc Tukey HSD, p < 0.05.

### A complete, annotated *H. erectifolium* reference genome

Motivated by the unique leaf morphology and drought response of *H. erectifolium*, we investigated its genomic and transcriptomic differences relative to the cultivated barley cultivar Morex. *H. erectifolium,* like barley, has a large diploid (2n=14) genome, reported with a haploid size of 2n = 9.49 ±0.05 pg, ∼4.6 Gbp by flow cytometry, while *H. vulgare* is 5.04 Gbp (Jakob et al., 2004; Doležel et al. 2018).

Using short-read sequencing, 10x Genomics Linked-Reads, we characterized the haploid genome of *H. erectifolium* via k-mer analysis, estimating a genome size of 4.4 Gbp, a repeat content of 75 %, and no detectable heterozygosity (Fig. **2a**). For a high-quality chromosome scale genome assembly, we used a combination of ONT long (321 Gb, 72× coverage, read N50 of 42 kb), and Illumina short (244.6 Gb, 55.6× coverage) read data, Bionano optical genome mapping (546 optical genome maps, map length of 3.8 Gbp, N50 of 20.1 Mbp) for hybrid scaffolding and Hi-C chromosomal conformation capture sequencing for pseudomolecule construction (72.2 Gb, 16.4× coverage) (Table **S1-S4**). The final assembly size was 3.94 Gbp with 3.85 Gbp (97.7 %) being anchored in seven pseudomolecules and 89 Mbp of unassigned contigs. (Fig. **2b,c**) (Table **S5**). The genome had a k-mer completeness and QV estimate score of 94 % and QV 34.6 and a BUSCO estimate of gene space completeness of 98.2 % (Single: 93.4 %, Duplicate:4.8 %, Poales database, n = 4,896) (Table **S5**) (Rhie et al., 2020; Manni et al., 2021).

**Fig. 2:**
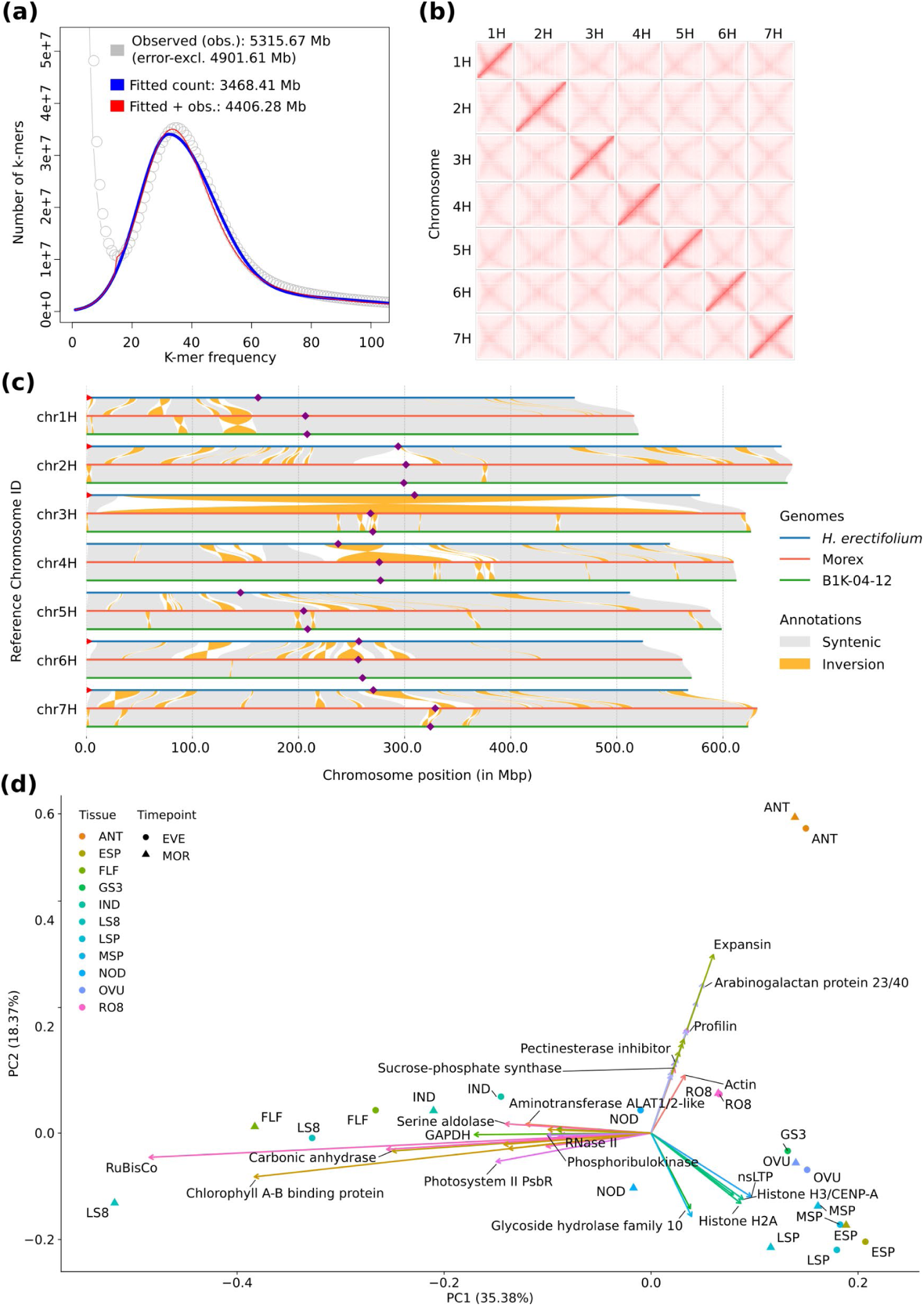
Genome characteristics, interchromosomal contact matrix, chromosomal synteny between genomes, and principal component analysis (PCA) of tissue-specific expression profiles. **(a)** K-mer spectra of 21-mers, calculated form short-read data, red line indicates fitted and observed 21-mers for genome size calculation. **(b)** Interchomosomal contact matrix of the seven assembled chromosomes. Pixel intensity represents Hi-C link counts normalized over a 1 Mb window on a logarithmic scale. **(c)** Whole chromosomal synteny between each respective chromosome in *H. erectifolium*, Morex, and B1K-04-12. Inversions smaller than 1 Mb were excluded for visualization. Purple diamonds indicate putative centromere location and red triangles indicate identified telomeric sequences. **(d)** PCA clustering of transcript abundance in 21 tissue-time specific samples and the trajectory of the top 20 highest loading genes for PC1 and PC2. Eleven individual tissues of which ten were sampled both in the morning (MOR, ZT 1-3) and evening (EVE, ZT 13-15). Opposite trajectories of vegetative versus reproductive tissues along PC1 and a third trajectory on PC2 contributed by anthers (ANT) and to a lesser extent, roots (RO8). Abbreviations: flag leaf (FLF), three-day-old germinating seeds (GS3), third internode (IND), fourth node (NOD), whole shoots, (LS8, 8 days post germination), whole roots (RO8, 8 days post germination), ovules (OVU), anthers (ANT), caryopses (CAR), developing shoot apical meristems: ESP (W3.0-4.5), MSP (W5.0-6.5), LSP (W7.0-8.0) (Waddington et al., 1987).

Taken together, we assembled a high-quality chromosomal-scale reference genome of *H. erectifolium* using ONT long reads and polished with Illumina short reads, attaining a consensus score of QV 34.6. The final genome consisted of 3.85 Gbp assigned to 7 pseudomolecules.

### Tissue-time specific long read transcriptome sequencing and gene annotation

Transcriptome profiling of 22 samples from 12 different tissues harvested at two diurnal timepoints, morning (MOR) and evening (EVE) yielded 3,645,621 full-length and non-chimeric reads used for annotation and analyses (Table **S6, S7**). Principal component analysis (PCA) grouped the MOR and EVE tissue pairs closely together, vegetative and reproductive samples differentiated along PC1 (35 %) with roots (RO8) and nodes (NOD) at the center. High expression of *PROLIFIN* and *EXPANSIN* genes separated anthers (ANT) from other reproductive samples (Fig. **2d**). When the caryopsis (CAR) was included, it accounted for 49 % of the PC1 variance, primarily due to high expression of multiple *GLIADIN* genes, which encode the major storage proteins in cereal grains (Fig. **S2**) (Anderson et al., 2012). The trajectory of developing shoot apical meristem (ESP, MSP, LSP) and ovules (OVU) were most influenced by histones (*H2A* and *H3/CENP-A*) and a non-specific lipid-transfer protein *(nsLTP*). Photosynthesis-related genes, particularly *CAB* (Chlorophyll a-b binding protein) and *RuBisCo* (Ribulose-1,5-bisphosphate carboxylase/oxygenase), were the main drivers distinguishing above-ground vegetative samples, including whole shoots (LS8) and flag leaves (FLF; Fig. **2d**, **S2**). We combined two structural gene annotation methods, where IsoSeq yielded 29,099 protein-coding genes (with 5’ and 3’ UTRs) and 8,993 putative lncRNAs, Helixer predicted 55,287 protein-coding genes, of which 20,875 fully and 1,029 partially overlapped with IsoSeq. We annotated a total of 71,475 genes, thereof 8,993 lncRNAs, and 157,682 isoforms. IsoSeq sequencing captured 98,238 unique protein-coding isoforms, with 3.4 isoforms per gene, 5.9 and 4.5 exons per mRNA and CDS, respectively. Mean gene, mRNA and CDS lengths were 5,835 bp, 3,808 bp and 826 bp, respectively, while lncRNA genes were considerably shorter at 1,857 bp, and only 1.8 isoforms and two exons per gene (Table **S8**).

In summary, with IsoSeq mRNA sequencing in 22 tissue samples, we could generate high-accuracy gene predictions with UTRs and accurate isoform information, as well as lncRNA predictions. Complemented with *ab initio* gene prediction, we identified and annotated a final set of 71,475 genes (thereof 8,993 lncRNAs) and 157,682 isoforms. Furthermore, we provided a comprehensive tissue and time of the day specific gene expression atlas in *H. erectifolium*.

### Comparative genome analysis between *H. erectifolium* and *H. vulgare*

We further characterized the genome of *H. erectifolium* in relation to the reference genomes of the spring barley cultivar Morex and the wild barley accession B1K-04-12 (Jayakodi et al., 2020; Mascher et al., 2021). We compared the chromosome organization and structure, repeat content and identified telomeric sequences, centromere locations, and transposable element (TE) composition. The assembly of telomeres and centromeres is one of the remaining challenges in gapless genome assemblies, due to the presence of long stretches of satellite repeats (Navrátilová et al., 2022). Here we report that the telomeric ends were only found on the short arms of five out of seven chromosomes (missing on 4H and 5H) in *H. erectifolium* (Fig. **2c**). No telomeric ends were found on long and short arms in either of the two barley genomes. We found *CRM* (homolog of *Cereba* in barley) enrichments in the pericentromeric regions spanning on average ∼37 Mbp per chromosome in *H. erectifolium,* and ∼100 intact *CRMs* per chromosome, but only ∼18 intact in barley (Fig. **2c**, **S3**) (Presting et al., 1998; Hudakova et al., 2001; Neumann et al., 2011). The *CRM* enrichment midpoints also colocalized with the regions identified in the Hi-C matrices (Fig. **1b**) (Navratilova et al., 2022).

Large-scale chromosomal synteny among *H. erectifolium*, Morex, and B1K-04-12 was overall conserved, although numerous small- to medium-scale structural rearrangements were detected. However, two notable large pericentric inversions were found, a near full chromosomal inversion on 3H that spanned 471 Mbp (81.4 %) in *H. erectifolium* and 570 Mbp (91.7 %) in Morex, and on 4H a pericentric inversion of 56 Mbp (10.2 %) in *H. erectifolium* and 109 Mbp (17.9 %) in Morex (Fig. **2c**). Neither inversion had previously been reported in barley (Mascher et al., 2024; Jayakodi et al., 2024). Between *H. erectifolium* and Morex, we found 59 inversions >1 Mbp, and 17 inversions >10 Mbp and thus more than between Morex and B1K-04-12, with 32 inversions >1 Mbp and one inversion >10 Mbp.

We performed *de novo* annotation of transposable elements (TEs) in the *H. erectifolium* genome and re-annotated TEs in the barley genomes Morex and B1K-04-12. TE content was relatively consistent across genomes, covering 85.7 % in *H. erectifolium*, 87.5 % in Morex, and 86.9 % in B1K-04-12. In *H. erectifolium*, the majority of repeats were LTR retrotransposons (68.2 %), while non-LTR elements included terminal inverted repeats (*TIRs*, 11.3 %) and Helitrons (5.3 %) (Table **S9**). Among LTRs, *Gypsy* elements were most abundant (20.1 %), followed by *Copia* (11.5 %), with 37.6 % classified as mixed or unclassified LTRs. Notably, CACTA transposons were enriched in *H. erectifolium* (7.2 %), but were less prevalent in Morex (1.3 %) and B1K-04-12 (2.2 %) (Table **S9**). Although overall LTR content was comparable across genomes, the number of intact LTRs varied by clade (Fig. **3b**). In *H. erectifolium*, *Retand* (39 %), *Angela* (23 %), *Tekay* (15 %), and *SIRE* (8 %) dominated, accounting for 85 % of all intact LTRs. In contrast, Morex and B1K-04-12 were primarily composed of *Angela* (39 %, 35 %), *Athila* (29 %, 35 %), *Tekay* (10 %, 8 %), and *SIRE* (7 %, 9 %), contributing 85 % and 87 % of their LTR content, respectively.

**Fig. 3:**
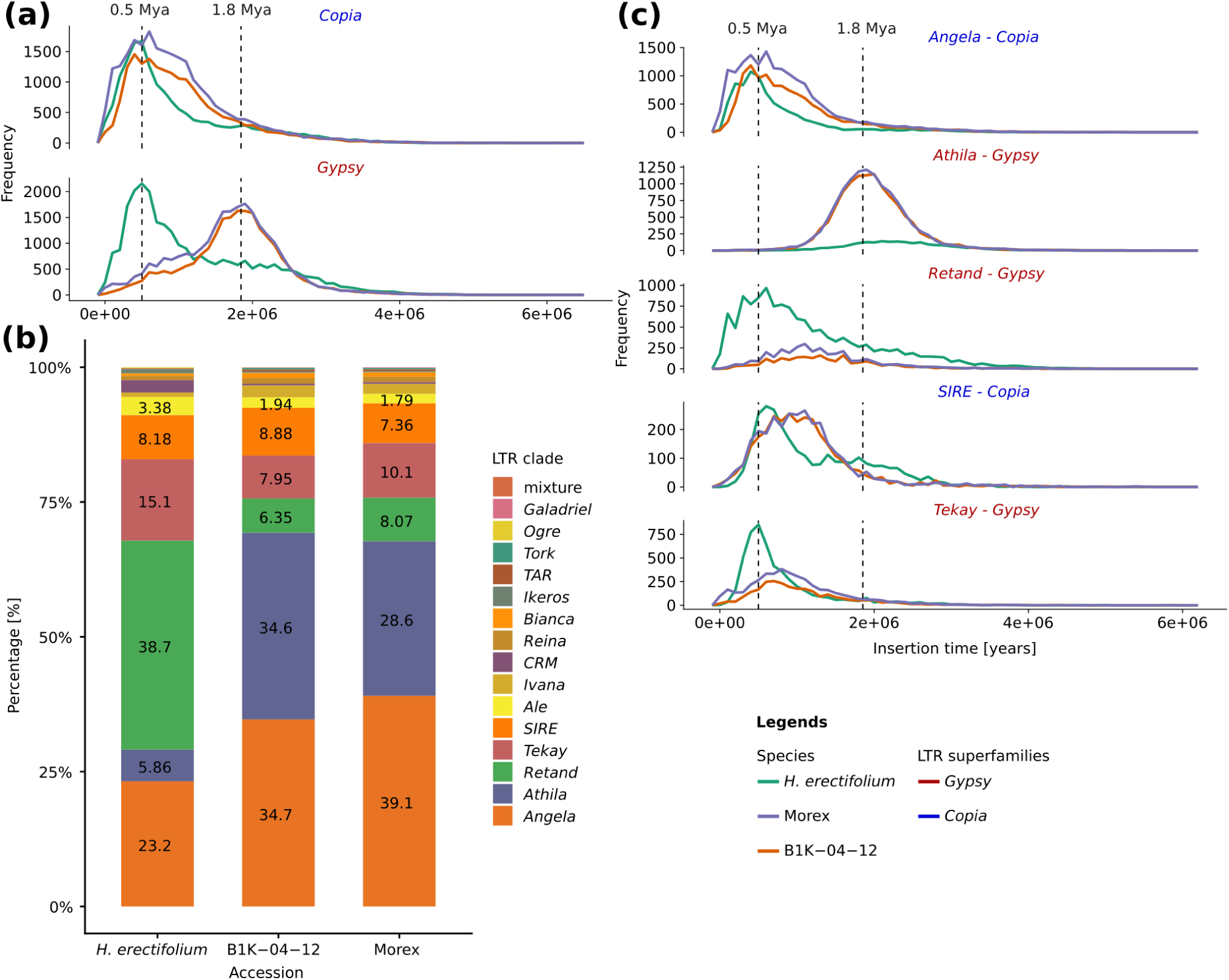
Transposable elements composition, long terminal repeat (LTR) retrotransposon profiles, and frequency of predicted insertion times. **(a)** Frequency plot of estimated LTR retrotransposon superfamily insertion times, *Gypsy* (red) and *Copia* (blue). Insertion times were calculated based on mutation rate in rice (1.3×10^-8^) of intact LTR retrotransposons in *H. erectifolium*, Morex, and B1K-04-12. Dotted lines mark LTR retrotransposon insertion burst peaks at 0.5 million years ago (Mya) and ∼1.8 Mya. **(b)** Composition landscape of intact LTR retrotransposons in *H. erectifolium*, Morex, and B1K-04-12. The percentage of LTR retrotransposons was normalized to the total of intact LTR retrotransposons within a genome. **(c)** Intact LTR retrotransposon insertion frequency over time by clade for the five most abundant clades; *Copia*: *Angela* and *SIRE* (blue), *Gypsy*: *Athila*, *Retand*, and *Tekay* (red). The black dotted lines mark 0.5 Mya and 1.8 Mya, respectively.

Insertion times of intact LTRs were estimated using a nucleotide substitution rate of 1.3×10⁻⁸ to infer their activation periods (Ma and Bennetzen, 2004; Ou and Jiang, 2018). We observed two distinct *Gypsy* insertion peaks: one at ∼0.5 Mya in *H. erectifolium*, and another at ∼1.8 Mya in Morex and B1K-04-12. *Copia* showed a concurrent insertion burst around 0.5 Mya in all three genomes during the Chibanian, coinciding with intensified and asynchronous glacial cycles (Fig. **3a**) (Sun et al., 2021). At this time, *Copia* insertions were predominantly driven by the *Angela* clade across all genomes, while *H. erectifolium* also showed lineage-specific bursts of the *Gypsy* clades *Tekay* and *Retand* (Fig. **3c**). The older burst at ∼1.8 Mya, unique to Morex and B1K-04-12, involved *Athila* insertions, corresponding with the Gelasian-Calabrian transition and early Northern Hemisphere glaciation (Gibbard et al., 2010; Cita et al., 2012). Recent *Copia* insertions were more frequent near chromosomal ends in all genomes, though also dispersed across chromosomes. Additionally, in *H. erectifolium*, *Gypsy* insertions exhibited a similar distribution pattern, recent (<0.6 Mya) and older (>0.6 Mya) (Fig. **S4**).

In summary, we identified numerous small- to medium-scale structural rearrangements between *H. erectifolium* and Morex, along with two large-scale pericentric inversions on chromosomes 3H and 4H. Despite similar TE amounts, each species displayed distinct LTR retrotransposon profiles, indicating independent TE-driven genome evolution. Additionally, we detected two major LTR insertion bursts across the *Hordeum* genus, occurring around 0.5 and 1.8 million years ago, coinciding with significant geological and climatic events.

### Genetic signatures of adaptive abiotic stress-related gene family expansions in *H. erectifolium*

We established hierarchical orthologous groups (HOGs) across the Poaceae family: *Sorghum bicolor Zea mays*, *Oryza sativa*, *Brachypodium distachyon*, *Triticum aestivum*, and *H. vulgare*, along with *Arabidopsis thaliana* as an outgroup, to explore unique or shared gene families, expansions, or contractions.

We identified 35,722 HOGs, 9,680 of which were shared across the Poaceae and Arabidopsis and 3,501 exclusive to Poaceae, while 1,472 HOGs were unique to *H. erectifolium* (Fig. **4a**). There were 7,182 genes in the 1,472 HOGs unique to *H.* erectifolium with Gene Ontology (GO) enrichments related to biological functions such as methylation, ribosome biogenesis, cell redox homeostasis, and signal peptide processing (Fig. **S5a**). Using a total of 18,869 HOGs with representative genes in at least two Poaceae species excluding *A. thaliana*, we identified 327 HOGs significant expanded or contracted in *H. erectifolium*, 288 expansions and 39 contractions (Fig. **S6**, Table **S10, S11**). The 3,968 genes from *H. erectifolium* belonging to the 327 HOGs were enriched for terms related to anatomical structure development, cell surface receptor signaling, and root development (Fig. **S5b**). Of those, we explored three expanded HOGs, with putative functions in abiotic stress resistance that had undergone expansions in *H. erectifolium* (Fig. **S7**, Table **S12**).

**Fig. 4:**
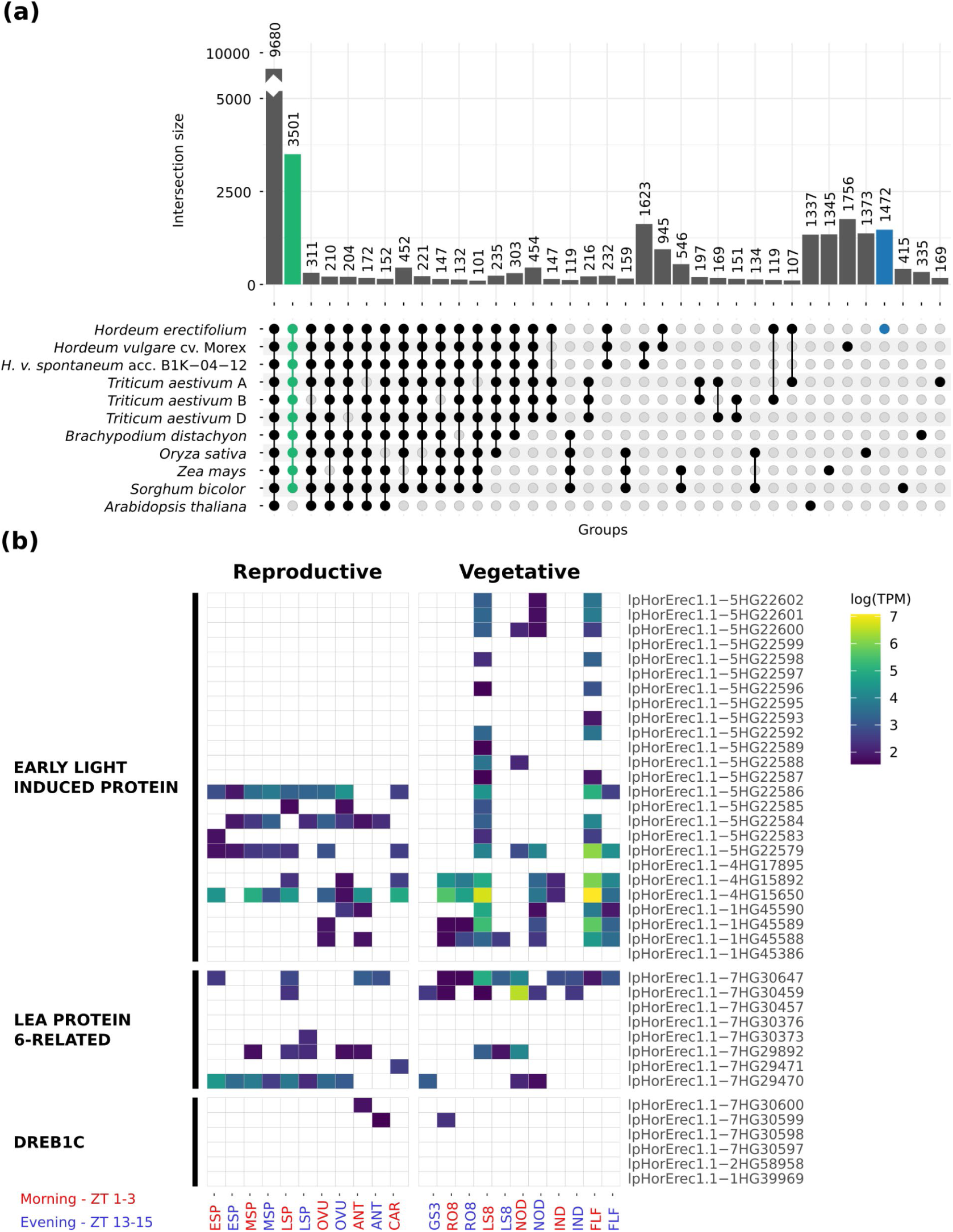
Shared and unique orthologs of *H. erectifolium* with Poaceae and *A. thaliana.* Tissue and time-specific expression of expanded gene families. **(a)** Shared and unique hierarchical phylogenetic orthologs (HOGs) between nine genomes of eight species; eight Poaceae species plus *A. thaliana*, the wheat (*T. aestivum*) subgenomes were separated; A, B, and D. Shared HOGs in Poaceae (green) and unique to *H. erectifolium* (blue) **(b)** Heatmap with expressed genes of expanded gene families with found in the 22 tissue-and-time specific samples; *EARLY LIGHT-INDUCED PROTEINS* (*ELIP*) (N0.HOG0000208), *LATE EMBRYOGENESIS ABUNDANT PROTEIN 6-RELATED* (*LEA PROTEIN 6-RELATED*) (N0.HOG0001214), and *DEHYDRATION-RESPONSIVE ELEMENT-BINDING PROTEIN 1C* (*DREB1C*) (N0.HOG0002366). Transcript abundance indicated as log(TPM), and timepoints: morning (red, MOR, ZT 1-3), and evening (blue, EVE, ZT 13-15). Tissues are divided in to reproductive: developing shoot apical meristems: ESP (W3.0-4.5), MSP (W5.0-6.5), LSP (W7.0-8.0), ovules (OVU), anthers (ANT), caryopses (CAR), and vegetative: three-day-old germinating seeds (GS3), whole roots (RO8, 8 days post germination), whole shoots, (LS8, 8 days post germination), fourth node (NOD), third internode (IND), flag leaf (FLF) (Waddington et al., 1987).

The *EARLY LIGHT-INDUCIBLE PROTEIN* (*ELIP*) gene family, which enhances desiccation tolerance by mitigating photooxidative damage (VanBuren et al., 2019), is expanded in *H. erectifolium* with 25 paralogs, compared to 20 in barley. The primary expansion occurred through tandem duplication on chromosome 5H, with 18 copies in *H. erectifolium* and 12 in barley. Of the 25 *ELIP* genes, 20 showed tissue- and time-specific expression: eleven were exclusively expressed in vegetative tissue, predominantly in leaves in the morning (LS8, FLF) whereas nine were expressed both in vegetative and reproductive tissue (Fig. **4b**, Table **S13**). We also observed an expansion of *LATE EMBRYOGENESIS ABUNDANT (LEA) PROTEIN 6-RELATED* genes (group 4 LEAs), which protect proteins and membranes during desiccation (Candat et al., 2014). *H. erectifolium* carried eight paralogs versus four in barley, with six showing expression in at least one tissue or time point (Fig. **4b**, Table **S13**). Similarly, the *DEHYDRATION-RESPONSIVE ELEMENT-BINDING PROTEIN 1C* (*DREB1C*, N0.HOG0002366) family underwent tandem duplication on chromosome 7H in *H. erectifolium*, resulting in four paralogs compared to two in barley; however, only two were expressed in the IsoSeq data, each in a single tissue (Fig. **4b**, Table **S13**).

In summary, gene sets unique to *H. erectifolium* were enriched for functions including methylation, ribosome biogenesis, cell redox homeostasis, and signal peptide processing. HOGs expanded in *H. erectifolium* were enriched for terms related to anatomical structure development, cell surface receptor signaling, and root development. Coupled with expansions of gene families associated with abiotic stress adaptation, i.e., the desiccation-related *ELIP*, *LEA*, and *DREB1C* genes, these enrichments indicate that *H. erectifolium* is genetically primed to tolerate environmental stress.

### Cross-species transcriptional drought response and recovery

To explore differences and similarities in the transcriptional responses to drought between *H. erectifolium* and barley cv. Morex, we set up a drydown drought and recovery experiment at the vegetative stage. Using a controlled drydown assay, we ensured that the reduction in leaf relative water content (RWC) was equal between plants of the two species in the control and water-limited conditions.

During drydown, both leaf RWC and soil field capacity (FC) decreased at the same rate in both species, with no significant differences between either species at each respective day-of-treatment (DOT) timepoint (Fig. **S8, S9**). Control samples were kept at 50 % soil FC while water was withheld for 6 days before rewatering, and samples were collected on days 2, 5, 6, and 7 of treatment. On the last day of drydown, DOT6, the average leaf RWC and soil FC for *H. erectifolium* were at 42 % ± 9 and 11 % ± 2, and for Morex at 51 % ± 9 and 13 % ± 0.6, respectively. Strong phenotypic differences at DOT5 and DOT6 were observed; leaves of *H. erectifolium* had rolled inward and remained erect, whereas Morex leaves wilted and required structural support (Fig. **S8b,c**). Both species recovered leaf RWC to control levels and regained leaf structure on DOT7, 24 hours after re-watering. No visible symptoms of senescence, i.e., no yellowing of leaves, were observed at DOT7 nor seven days after re-watering, DOT13.

In total, 64 leaf samples from the two species, four time points and two treatments with four biological replicates per sampling point, were RNA-sequenced with an average of 21 million reads per sample (Table **S14**). The RNAseq data were mapped to their respective genomes with STAR and quantified with featureCounts (Table **S15**) (Dobin et al., 2013; Liao et al., 2014). We employed two methods for transcriptomic analysis: maSigPro for time-course analysis, taking into account day of sampling and treatment variables (Nueda et al., 2014), and edgeR for pairwise expression comparison between control and drydown for each time point separately, to also identify transiently expressed genes (Chen et al., 2025).

With both methods, we identified 15,036 and 13,119 differentially expressed genes (DEGs) in *H. erectifolium* and Morex, respectively (Table **S16-S18**). The time-course analysis identified 7,941 DEGs in *H. erectifolium* and 7,382 DEGs in Morex (Fig. **S10**). In addition, we detected 7,095 DEGs in *H. erectifolium* and 5,737 DEGs in Morex, which were expressed only at specific DOTs and not identified in the time-course analysis.

To compare DEGs between the two species, we focused on their shared single-copy orthologs (SCO). Since transient DEGs showed a low overlap between species, we only included DEGs identified by the timecourse analysis. Between *H. erectifolium* and Morex, we identified 19,204 SCO, of which 8,451 were differentially expressed in at least one species, thereof 5,487 in *H. erectifolium* and 5,743 in Morex, with an overlap of 2,799 (33 %) DEGs detected in both *H. erectifolium* and Morex (Fig. **5a**, Table **S19**).

**Fig. 5:**
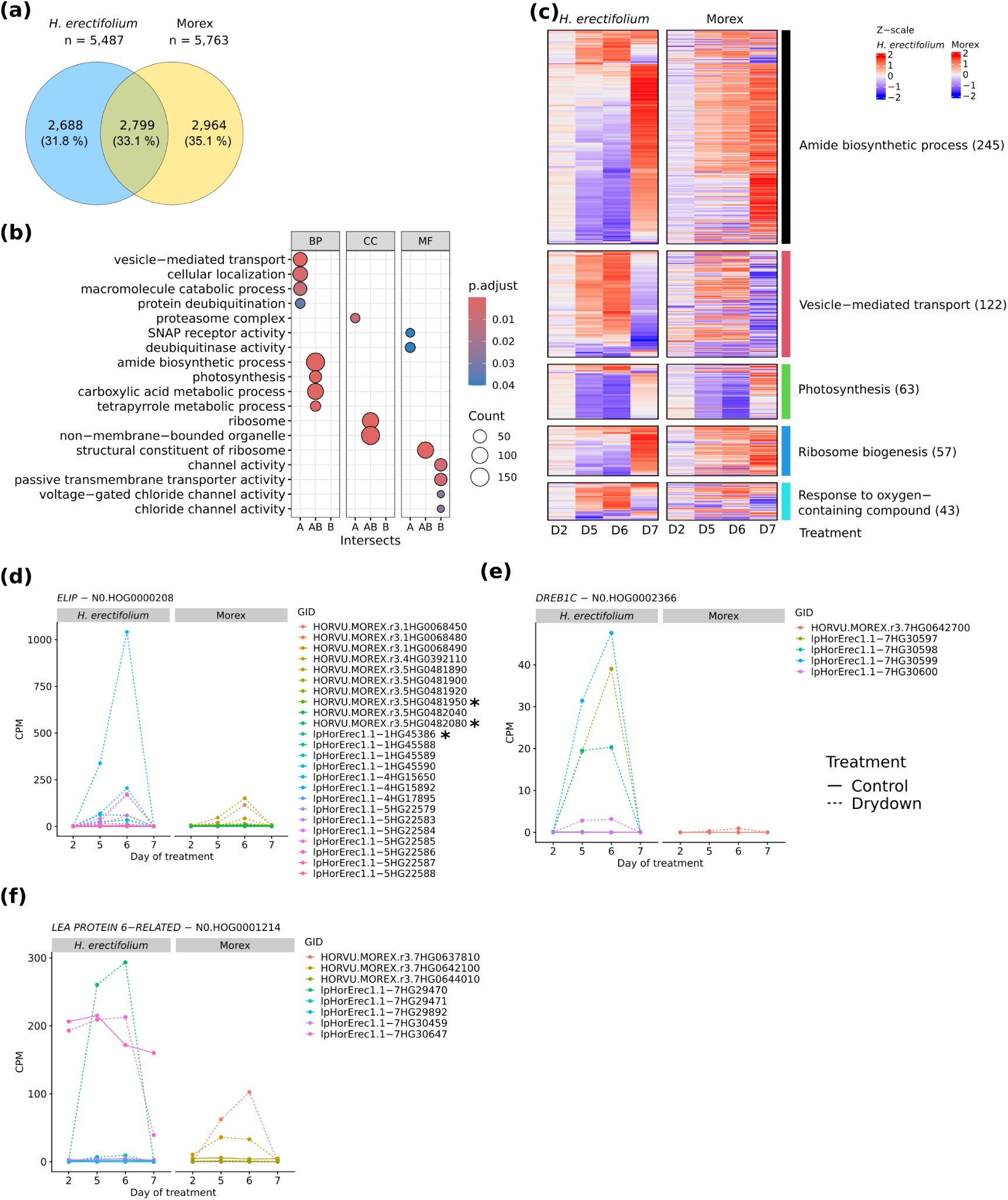
Differentially expressed genes (DEG) during drydown and recovery in *H. erectifolium* and Morex. **(a)** Total number of shared and exclusive single-copy ortholog (SCO) genes differentially expressed between *H. erectifolium* and Morex. **(b)** GO enrichment of shared and exclusive SCO DEGs: exclusive to *H. erectifolium* (A), exclusive to Morex (B), and interspecies overlap (AB). GO categories: biological processes (BP), cellular compartments (CC), and molecular functions (MF). **(c)** Five groups of the enriched biological processes terms in SCO: amide biosynthetic process, vesicle-mediated transport, photosynthesis, ribosome biogenesis, and response to oxygen – containing compound, total number of genes shown in brackets. The heatmap shows scaled log(fold-change), and gene clustering was based on *H. erectifolium* transcripts with the corresponding SCO in Morex fixed to the same location in the heatmap. Second day-of-treatment (DOT) (D2); fifth DOT (D5); D6, sixth DOT (D6); seventh DOT (D7, recovery). Gene expression of expanded hierarchical phylogenetic orthologs (HOG) gene families in *H. erectifolium,* and Morex genes found in the same HOG, in response to drydown and recovery. **(d)** *EARLY LIGHT-INDUCED PROTEINS* (*ELIP*) (N0.HOG0000208), **(e)** *LATE EMBRYOGENESIS ABUNDANT PROTEIN 6-RELATED* (*LEA PROTEIN 6-RELATED*) (N0.HOG0001214), **(f)** *DEHYDRATION-RESPONSIVE ELEMENT-BINDING PROTEIN 1C* (*DREB1C*) (N0.HOG0002366). Graphs show normalized counts per million (CPM) values for control and drydown. All genes, except those marked with asterisk, were differentially regulated in response to drydown or recovery.

DEGs shared by both species were enriched for photosynthesis, translation (ribosome), amide biosynthesis, carboxylic acid and tetrapyrrole metabolism, suggesting that both species adapt metabolically and coordinate energy production, carbon and nitrogen assimilation, and oxidative stress management in response to water limitations (Fig. **5b**). However, while both species downregulated genes with functions in photosynthesis during water limitation, they exhibited markedly different responses in genes related to biosynthesis and translation. Notably, differentially expressed genes (DEGs) associated with biosynthesis (amide biosynthetic processes) and translation (ribosome-related processes) were strongly downregulated in *H. erectifolium*, whereas they were upregulated in Morex at DOT5 and DOT6 (Fig. **5c**). In *H. erectifolium*, the downregulation of these DEGs indicated a repression of biosynthesis and translational activity in response to drought. The downregulation of amide biosynthesis, i.e., decreased transcript levels of glutamine and asparagine synthase genes (Table **S17, S18**), likely reflected a metabolic shift towards energy conservation to prioritize survival over growth. The concurrent upregulation of ubiquitination, macromolecule catabolism, and vesicle-mediated transport specifically in *H. erectifolium* indicated mobilization of proteins for recycling or stress adaptation and active degradation of damaged proteins and cellular components. In contrast, Morex upregulated genes involved in amide biosynthesis and translation, potentially as part of a protective response to counteract ROS-induced protein denaturation and to replace damaged proteins to maintain cell functions. The transcriptional responses in Morex were thus reminiscent of proteostasis responses, which commonly lead to massive transcriptional and translational upregulation of protective proteins (Mittler et al., 2012). The Morex-specific enrichment of DEGs with functions in transmembrane transport and chloride channel activities suggested that Morex maintained ion and osmotic balance to support active metabolism under water limitations. Morex thus invested in maintaining cellular functions and supporting future recovery rather than shutting down. In line with this, at recovery (DOT7), transcripts related to biosynthesis and translation were strongly induced in *H. erectifolium* but not strongly altered compared to DOT5 and 6 in Morex (Fig. **5c**).

Given the different transcriptional responses to water limitation between both genotypes, we further explored the expression of the three expanded desiccation-tolerance gene families, encoding *DREB*, *ELIP*, and *LEA* proteins, which function in cellular protection and stabilization, particularly during dehydration or oxidative stress. Among the *ELIP* (N0.HOG0000208) genes, 13 of 25, and eight of 20 were differentially expressed during desiccation in *H. erectifolium* and Morex, respectively (Fig. **5d**, **S11a**). We observed a strong upregulation of the four *DREB1C* genes (N0.HOG0002366) in *H. erectifolium* compared to only a mild upregulation of one *DREB1C* copy in Morex (Fig. **5f**, **S11b**)*. DREB* transcription factors act as upstream regulators that activate *LEA* genes, which were also more strongly expressed in *H. erectifolium* than in Morex (Sakuma et al., 2006). Of the *LEA* (N0.HOG0001214) genes, five of the eight copies were upregulated in *H. erectifolium*, whereas only three of four copies were upregulated in Morex in response to drought (Fig. **5e**, **S11c**). One *LEA* (lpHorErec1.1-7HG30647) had a high constitutive expression under control and drought in *H. erectifolium* and was also expressed in nearly all vegetative tissues (Fig. **4b**).

Taken together, the differential regulation of amide biosynthesis and translation in *H. erectifolium* and Morex suggested contrasting metabolic responses to water limitations. Morex upregulated both pathways, thereby maintaining active nitrogen metabolism and detoxification and proteostasis. In contrast, *H. erectifolium* downregulated both pathways, indicative of a conservative, survival-focused strategy, minimizing energy expenditure and growth processes to preserve cellular integrity during drought. The expanded and highly inducible *DREB*, *LEA*, and *ELIP* families suggested that *H. erectifolium* is genetically primed for drought, enabling strong protective responses with low metabolic cost.

## Discussion

### The unique leaf morphology of *H. erectifolium*

*H. erectifolium* displayed distinct leaf morphological traits indicative of drought adaptation, including dense trichome coverage, high vein density with a near-equal major-to-minor vein ratio, and a thick cuticular wax layer, all features of xeromorphic leaf traits (Shields, 1950). The prominently ribbed adaxial surface bore dense short trichomes and longer trichomes along major veins, features involved in microclimate stabilization, dew capture, and enhanced structural rigidity (Liakoura et al., 1997; Galdon-Armero et al., 2018). A compact, uniform epidermal cell layer and substantial wax coating likely strengthened the transpiration barrier, UV protection and prevented biotic intrusion (Kasapligil, 1961; Xue et al., 2017). Major veins were reinforced with extensive bundle sheath extensions (BSE), sclerenchymatous tissue causing increased mesophyll compartmentalization which has been associated with enhanced light penetration, photosynthetic efficiency, and water-use optimization through more responsive BSE-connected stomatal regulation (Barbosa et al., 2019). *H. erectifolium* and Morex both maintained comparable leaf RWC levels as soil moisture declined during drydown. However, while Morex leaves wilted and required external support, *H. erectifolium* leaves rolled tightly and remained erect (Fig. **1**, **S8**). Leaf rolling under water deficit reduces exposed surface area, minimizing transpiration, photodamage, and heat stress (Kadioglu et al., 2012). Although bulliform cells are typically associated with this response, they were largely absent in *H. erectifolium* (Cal et al., 2019; Zhu et al., 2022). We propose that leaf rolling in this species is facilitated by the ribbed structure, numerous major veins with extensive BSE and sclerenchyma, and a compact epidermal cell architecture (Guo et al., 2024). The distinctive morphology of *H. erectifolium* was mirrored at the genomic level by an enrichment of expanded gene families involved in anatomical structure development.

Triticeae genomes are typically large (∼5 Gbp per genome or subgenome) and consist of up to 90 % repetitive elements, primarily transposable elements (Middleton et al., 2013; Winterfeld et al., 2025). While the total proportion of repetitive content was relatively stable across species (85.7–87.5 %), we observed notable differences in LTR clade composition and insertion histories. *H. erectifolium* showed a five-fold enrichment of CACTA transposons compared to barley, elements that influence genome restructuring and gene evolution (Lisch, 2013; Catoni et al., 2019; Liu et al., 2020). We discovered two LTR retrotransposon insertion bursts at ∼0.5 Mya and ∼1.8 Mya, which coincide with known geological events. The Chibanian stage (∼0.5 Mya), characterized by intense, globally asynchronous glacial cycles, and the Calabrian stage (Early Pleistocene, ∼1.8 Mya), marked by the onset of Northern Hemisphere glaciation affecting Eurasian species (Cita et al., 2012; Sun et al., 2021). There was a synchronous peak at ∼0.5 Mya, where *Angela* (*Copia*) was active in both species, while the *Gypsy* families *Retand* and *Tekay* were specific to *H. erectifolium*. The second unilateral peak at ∼1.8 Mya, which was observed only in barley, was exclusively associated with *Athila* (*Gypsy*) (Fig. **4**). LTR insertion bursts 1.8 Mya were also detected in two other Mediterranean species, wheat and Brachypodium distachyon (Wicker et al., 2018; Stritt et al., 2020), suggesting that Eurasian species experienced more severe glaciation and biome turnover than South American species (Vuilleumier 1971; Connor 2009; Gibbard et al., 2010). Accordingly, phylogenetic and biogeographic studies suggested high extinction rates for the Eurasian *Hordeum* species during the Early Pleistocene, while high speciation rates were maintained in the South American clade (Jacob and Blattner, 2006).

The higher copy number of desiccation-related genes and the distinct transcriptome response to water limitation indicated that *H. erectifolium* and Morex diverge not only in their morphological but also in their molecular responses to stress. Gene family expansion through tandem duplication can enable faster transcriptional activation and greater dosage effects, providing an adaptive advantage during rapid declines in relative water content (Dassanayake et al., 2011; Wu et al., 2012). Large tandem arrays of *ELIP*s, which stabilize and protect photosynthetic components during dehydration and enhance thermal energy dissipation, have been repeatedly associated with independent origins of desiccation tolerance across diverse plant lineages (VanBuren et al., 2019; Pardo et al. 2020; Marks et al., 2024). Similarly, the observed expansion of *DREB1C* and *LEA* gene families in *H. erectifolium* is consistent with a genetic architecture that enhances resilience across multiple stresses, including drought, salinity, and cold (Hernández-Sánchez et al., 2022; Wang et al., 2022). Such expansions likely genetically primed *H. erectifolium* for rapid adaptation to fluctuating water availability, offering a selective advantage in environments where desiccation events are frequent and unpredictable. This genetic priming for rapid stress response is reminiscent of mechanisms seen in resurrection plants, where tandem gene duplications of protective proteins underpin efficient tolerance with minimal metabolic burden (VanBuren et al., 2019; Pardo et al. 2020).

The drydown experiment corroborates this hypothesis, showing fast and strong activation of all three gene families followed by rapid deactivation upon rehydration, a pattern indicative of efficient regulatory control. Furthermore, the constitutive high expression of one *LEA* copy may have conferred an immediate protective buffer at the onset of dehydration stress, minimizing cellular damage during early water loss (Cheng et al., 2002; Liu et al., 2009).

The transcriptional responses of *H. erectifolium* and Morex to water limitation suggested contrasting drought-adaptation strategies: metabolic downregulation and survival prioritization in *H. erectifolium* versus maintenance of metabolic activity and competitiveness in Morex (Skirycz et al., 2011; Claeys & Inzé, 2013). Both genotypes showed enrichment of DEGs related to photosynthesis, translation, and nitrogen metabolism, but their regulation differed markedly. In *H. erectifolium*, strong repression of amide biosynthesis and ribosome-related processes, including reduced expression of glutamine and asparagine synthetases, indicates energy conservation through suppression of growth and nitrogen assimilation (Diaz et al., 2010, Nagy et al., 2013). Concurrent induction of ubiquitination and macromolecule catabolism suggests active recycling of proteins and degradation of damaged components to protect cellular integrity (Skirycz & Inzé, 2010; Eckardt et al., 2024). Furthermore, *H. erectifolium* showed unique activation of pathways linked to cellular localization, vesicle-mediated transport, and oxidative stress mitigation, including ROS metabolism, which together indicate pre-emptive preparation for drought (Mazel et al., 2004; Jarzyniak & Jasiński, 2014; Noctor et al., 2014). By contrast, Morex upregulated biosynthesis and translation, consistent with a strategy of sustaining metabolic activity and proteostasis to replace damaged proteins and maintain cellular function (Mittler et al., 2012). The enrichment of transmembrane transport and chloride channel activity indicates active osmotic regulation, supporting growth and recovery under moderate stress (Nieves-Cordones et al., 2019; Franco-Navarro et al., 2021). The absence of strong post-drought induction of biosynthetic and translational processes in Morex at recovery, compared to the rapid reactivation seen in *H. erectifolium*, further highlights differences in resource allocation and stress recovery dynamics.

Overall, *H. erectifolium* appears genetically primed for survival under severe drought through gene family expansion and rapid protective responses, efficient shutdown of growth-related metabolism, and rapid recovery. Such a strategy is likely advantageous in habitats with frequent and severe desiccation events, favoring survival over productivity. Morex, in contrast, invests in maintaining metabolic activity during drought, which may favor productivity and recovery potential under mild stress but is energetically costly under prolonged or severe water deficits (Skirycz et al., 2011).

Combining both strategies, stress avoidance through active metabolism with drought tolerance via growth suppression, is a central challenge for breeding crops that can both survive severe drought and maintain yield under moderate stress. The exploration of desiccation-tolerant crop relatives might provide novel genetic insights and tools, such as stress-inducible protective pathways (e.g., *DREB*, *LEA*, *ELIP*) and stress-responsive promoters to activate key protective genes while minimizing growth penalties under transient or moderate stress (Nelson et al., 2007; Peleg et al., 2011).

## Acknowledgements

We greatly thank Rebekka Schüller and Nina Döring for excellent technical assistance, and Dominik Brilhaus for data management support. This work was supported by the by the DFG Research Infrastructure NGS_CC (West German Genome Center (WGGC), project 407493903) as part of the Next Generation Sequencing Competence Network (project 423957469). Computational infrastructure and support were provided by the Centre for Information and Media Technology at Heinrich Heine University Düsseldorf. This work was funded by the European Research Council (ERC) under the European Union’s Horizon Europe research and innovation programme (PERLIFE, No. 101002085 to MvK and TRANSFER, No. 949873 to MM), the Deutsche Forschungsgemeinschaft (DFG) under Germany’s Excellence Strategy—EXC-2048/1—Project ID: 390686111, and the Collaborative Research Centre/Transregio (TRR 341, Project ID: 456082119), and the GRK 2064: Water use efficiency and drought stress responses: From Arabidopsis to Barley, Project ID: 252965955. H. Šimková and Z. Tulpová were supported from the project TANGENC, reg. no. CZ.02.01.01/00/22_008/0004581 of the ERDF Programme Johannes Amos Comenius.

## Author contributions

EBH and MvK conceived and designed the project and experiments. EBH performed all plant growth, DNA extractions, data processing, genome assembly, experiments, and analysis, with the following exceptions: HŠ, HT, and ZT made the optical genome map and did the hybrid scaffolding. MA provided the carbon-nitrogen elemental and Specific Leaf-Area measurements. TR performed RNA extractions for sequencing. MM and JF helped with the chromosome-scale genome sequence assembly. EBH wrote the manuscript with the help of MvK.

## Data availability

The data that support the findings of this study will be made openly available in DataPLANT and new sequence data generated in the study will be deposited at the European Nucleotide Archive (ENA).

## Supporting Information – Supplemental Tables

**Table S1:** Oxford Nanopore Technologies long read sequencing data.

**Table S2:** 10x Genomics linked short read Illumina sequencing data.

**Table S3:** Hi-C chromosomal conformation capture Illumina sequencing data.

**Table S4:** Optical genome maps - Bionano.

**Table S5:** Genome assembly metrics, completeness and quality score. From draft assembly, to finally pseudomolecules.

**Table S6:** Overview of PacBio IsoSeq full-length capture transcript RNA sequencing of tissue-time specific samples.

**Table S7:** Gene expression in the 22 tissue and time specific samples, read counts TPM normalized per sample with IsoQuant.

**Table S8:** Summary of gene annotation, combined from evidence based gene prediction and *ab initio,* and lncRNA.

**Table S9:** Transposable element annotation with EDTA in *H. erectifolium* acc. NGB6816, *H.v.* cv. Morex, and *H.v. spontaneum* acc. B1K-04-12.

**Table S10:** Total number of expanded or contracted hierarchical phylogenetic orthologs (HOG) across eight genomes from seven species.

**Table S11:** All significantly expanded or contracted hierarchical phylogenetic orthologs (HOG) in *H. erectifolium*.

**Table S12:** Desiccation related expanded gene families *ELIP*, *DREB1C*, and *LEA*.

**Table S13:** Tissue and time specific expression of the expanded gene families: *ELIP*, *DREB1C*, and *LEA* genes in the 22 IsoSeq samples.

**Table S14:** Transcriptome sequencing with Illumina PE150 of 64 leaf samples from *H. erectifolium* and Morex during drydown experiment.

**Table S15:** RNAseq mapping rates of each sample to their respective genome.

**Table S16:** Total number of significantly expressed genes found in *H. erectifolium* and Morex during drydown experiment.

**Table S17:** All expressed genes found in *H. erectifolium* during drydown experiment.

**Table S18:** All expressed genes found in Morex during drydown experiment.

**Table S19:** Cross species comparison of significant differentially expressed single copy orthologs genes using between *H. erectifolium* and Morex.

## Supporting Information - Supplemental figures

**Fig. S1:** Quantitative leaf characteristics of the leaf below flag leaf (LBF) in *H. erectifolium*, cultivated (Morex) and wild barley.

**Fig. S2:** Principal component analysis (PCA) of tissue-specific expression profiles.

**Fig. S3:** Putative centromere locations and pericentric sizes in *H. erectifolium*, Morex, and B1K-04-12.

**Fig. S4:** *Copia* and *Gypsy* LTR retrotransposon insertions over time in chromosomes 2H and 7H.

**Fig. S5:** Enriched biological pathways found in hierarchical phylogenetic orthologs.

**Fig. S6:** Summary of expanded and contracted gene families found by CAFE5.

**Fig. S7:** Significantly expanded gene families found in *H. erectifolium* related to drydown adaptation.

**Fig. S8:** Leaf relative water content and plant morphology during drydown and recovery.

**Fig. S9:** Soil field capacity and fresh weight biomass during drydown and recovery.

**Fig. S10:** Time-course analysis of transcriptome changes over time in response to drydown.

**Fig. S11:** Gene expression of expanded hierarchical phylogenetic orthologs gene families in response to drydown and recovery.

## References

Anderson OD, Dong L, Huo N, Gu YQ. 2012. A New Class of Wheat Gliadin Genes and Proteins. PLOS ONE 7: e52139.

Aury J-M, Istace B. 2021. Hapo-G, haplotype-aware polishing of genome assemblies with accurate reads. NAR Genomics and Bioinformatics 3: lqab034.

Badr A, M K, Sch R, Rabey HE, Effgen S, Ibrahim HH, Pozzi C, Rohde W, Salamini F. 2000. On the Origin and Domestication History of Barley (*Hordeum vulgare*). Molecular Biology and Evolution 17: 499–510.

Baker L, Grewal S, Yang C, Hubbart-Edwards S, Scholefield D, Ashling S, Burridge AJ, Przewieslik-Allen AM, Wilkinson PA, King IP, et al. 2020. Exploiting the genome of Thinopyrum elongatum to expand the gene pool of hexaploid wheat. Theoretical and Applied Genetics 133: 2213– 2226.

Barbosa MAM, Chitwood DH, Azevedo AA, Araújo WL, Ribeiro DM, Peres LEP, Martins SCV, Zsögön A. 2019. Bundle sheath extensions affect leaf structural and physiological plasticity in response to irradiance. Plant, Cell & Environment 42: 1575–1589.

Blattner FR. 2018. Taxonomy of the Genus Hordeum and Barley (Hordeum vulgare). In: Stein N, Muehlbauer GJ, eds. The Barley Genome. Cham: Springer International Publishing, 11–23.

Blum M, Chang H-Y, Chuguransky S, Grego T, Kandasaamy S, Mitchell A, Nuka G, Paysan-Lafosse T, Qureshi M, Raj S, et al. 2021. The InterPro protein families and domains database: 20 years on. Nucleic Acids Research 49: D344–D354.

Bohra A, Kilian B, Sivasankar S, Caccamo M, Mba C, McCouch SR, Varshney RK. 2022. Reap the crop wild relatives for breeding future crops. Trends in Biotechnology 40: 412–431.

Bolger M, Schwacke R, Usadel B. 2021. MapMan Visualization of RNA-Seq Data Using Mercator4 Functional Annotations. In: Dobnik D, Gruden K, Ramšak Ž, Coll A, eds. Solanum tuberosum: Methods and Protocols. New York, NY: Springer US, 195–212.

Bothmer R von, Jacobsen N, Baden C, Jorgensen RB, Linde-Laursen I. 1995. An ecogeographical study of the genus Hordeum (2nd edition).

Bothmer R von, Jacobsen N, Jørgensen RB. 1985. Two New American Species of Hordeum (Poaceae). Willdenowia 15: 85–90.

Bothmer R von, Komatsuda T. 2010. Barley Origin and Related Species. In: Ullrich SE, ed. Barley. Wiley, 14–62.

Brassac J, Blattner FR. 2015. Species-Level Phylogeny and Polyploid Relationships in Hordeum (Poaceae) Inferred by Next-Generation Sequencing and In Silico Cloning of Multiple Nuclear Loci. Systematic Biology 64: 792–808.

Brozynska M, Furtado A, Henry RJ. 2016. Genomics of crop wild relatives: expanding the gene pool for crop improvement. Plant Biotechnology Journal 14: 1070–1085.

Buchfink B, Reuter K, Drost H-G. 2021. Sensitive protein alignments at tree-of-life scale using DIAMOND. Nature Methods 18: 366–368.

Buckley TN, Sack L, Gilbert ME. 2011. The Role of Bundle Sheath Extensions and Life Form in Stomatal Responses to Leaf Water Status. Plant Physiology 156: 962–973.

Cai W, McPhaden MJ, Grimm AM, Rodrigues RR, Taschetto AS, Garreaud RD, Dewitte B, Poveda G, Ham Y-G, Santoso A, et al. 2020. Climate impacts of the El Niño–Southern Oscillation on South America. Nature Reviews Earth & Environment 1: 215–231.

Cal AJ, Sanciangco M, Rebolledo MC, Luquet D, Torres RO, McNally KL, Henry A. 2019. Leaf morphology, rather than plant water status, underlies genetic variation of rice leaf rolling under drought. Plant, Cell & Environment 42: 1532–1544.

Camacho C, Coulouris G, Avagyan V, Ma N, Papadopoulos J, Bealer K, Madden TL. 2009. BLAST+: architecture and applications. BMC Bioinformatics 10: 1–9.

Candat A, Paszkiewicz G, Neveu M, Gautier R, Logan DC, Avelange-Macherel M-H, Macherel D. 2014. The Ubiquitous Distribution of Late Embryogenesis Abundant Proteins across Cell Compartments in Arabidopsis Offers Tailored Protection against Abiotic Stress. The Plant Cell 26: 3148–3166.

Catoni M, Jonesman T, Cerruti E, Paszkowski J. 2019. Mobilization of Pack-CACTA transposons in Arabidopsis suggests the mechanism of gene shuffling. Nucleic Acids Research 47: 1311–1320.

Chen Y, Chen L, Lun ATL, Baldoni PL, Smyth GK. 2025. edgeR v4: powerful differential analysis of sequencing data with expanded functionality and improved support for small counts and larger datasets. Nucleic Acids Research 53: gkaf018.

Cheng Z, Targolli J, Huang X, Wu R. 2002. Wheat LEA genes, PMA80 and PMA1959, enhance dehydration tolerance of transgenic rice (Oryza sativa L.). Molecular Breeding 10: 71–82.

Cita MB, Gibbard PL, Head MJ, Alloway B, Beu AG, Coltorti M, Gibbard PL, Hall VM, Head MJ, Jiaqi L, et al. 2012. Formal Ratification of the GSSP for the Base of the Calabrian Stage (Second Stage of the Pleistocene Series, Quaternary System). Episodes 35: 388–397.

Claeys H, Inzé D. 2013. The Agony of Choice: How Plants Balance Growth and Survival under Water-Limiting Conditions. Plant Physiology 162: 1768–1779.

Connor SE. 2009. Human impact – the last nail in the coffin for ancient plants? Journal of Biogeography 36: 485–486.

Danecek P, Bonfield JK, Liddle J, Marshall J, Ohan V, Pollard MO, Whitwham A, Keane T, McCarthy SA, Davies RM, et al. 2021. Twelve years of SAMtools and BCFtools. GigaScience 10: giab008.

Dassanayake M, Oh D, Hong H, Bohnert HJ, Cheeseman JM. 2011. Transcription strength and halophytic lifestyle. Trends in Plant Science 16: 1–3.

Díaz P, Betti M, Sánchez DH, Udvardi MK, Monza J, Márquez AJ. 2010. Deficiency in plastidic glutamine synthetase alters proline metabolism and transcriptomic response in Lotus japonicus under drought stress. New Phytologist 188: 1001–1013.

Dobin A, Davis CA, Schlesinger F, Drenkow J, Zaleski C, Jha S, Batut P, Chaisson M, Gingeras TR. 2013. STAR: ultrafast universal RNA-seq aligner. Bioinformatics 29: 15–21.

Doležel J, Čížková J, Šimková H, Bartoš J. 2018. One Major Challenge of Sequencing Large Plant Genomes Is to Know How Big They Really Are. International Journal of Molecular Sciences 19: 3554.

Duc G, Agrama H, Bao S, Berger J, Bourion V, De Ron Antonio M., Gowda Cholenahalli L. L., Mikic Aleksandar, Millot Dominique, Singh Karam B., et al. 2015. Breeding Annual Grain Legumes for Sustainable Agriculture: New Methods to Approach Complex Traits and Target New Cultivar Ideotypes. Critical Reviews in Plant Sciences 34: 381–411.

Eckardt NA, Avin-Wittenberg T, Bassham DC, Chen P, Chen Q, Fang J, Genschik P, Ghifari AS, Guercio AM, Gibbs DJ, et al. 2024. The lowdown on breakdown: Open questions in plant proteolysis. The Plant Cell 36: 2931–2975.

Emms DM, Kelly S. 2019. OrthoFinder: phylogenetic orthology inference for comparative genomics. Genome Biology 20: 1–14.

Farooq M, Frei M, Zeibig F, Pantha S, Özkan H, Kilian B, Siddique KHM. 2025. Back into the Wild: Harnessing the Power of Wheat Wild Relatives for Future Crop and Food Security. Journal of Experimental Botany: eraf141.

Feuillet C, Langridge P, Waugh R. 2008. Cereal breeding takes a walk on the wild side. Trends in Genetics 24: 24–32.

Franco-Navarro JD, Díaz-Rueda P, Rivero-Núñez CM, Brumós J, Rubio-Casal AE, de Cires A, Colmenero-Flores JM, Rosales MA. 2021. Chloride nutrition improves drought resistance by enhancing water deficit avoidance and tolerance mechanisms. Journal of Experimental Botany 72: 5246–5261.

Galdon-Armero J, Fullana-Pericas M, Mulet PA, Conesa MA, Martin C, Galmes J. 2018. The ratio of trichomes to stomata is associated with water use efficiency in Solanum lycopersicum (tomato). The Plant Journal 96: 607–619.

Gao M, Hao Z, Ning Y, He Z. 2024. Revisiting growth–defence trade-offs and breeding strategies in crops. Plant Biotechnology Journal 22: 1198–1205.

Gibbard PL, Head MJ, Walker MJC, The Subcommission On Quaternary Stratigraphy. 2010. Formal ratification of the Quaternary System/Period and the Pleistocene Series/Epoch with a base at 2.58 Ma. Journal of Quaternary Science 25: 96–102.

Goel M, Schneeberger K. 2022. plotsr: visualizing structural similarities and rearrangements between multiple genomes. Bioinformatics 38: 2922–2926.

Goel M, Sun H, Jiao W-B, Schneeberger K. 2019. SyRI: finding genomic rearrangements and local sequence differences from whole-genome assemblies. Genome Biology 20: 277.

Goodstein DM, Shu S, Howson R, Neupane R, Hayes RD, Fazo J, Mitros T, Dirks W, Hellsten U, Putnam N, et al. 2012. Phytozome: a comparative platform for green plant genomics. Nucleic Acids Research 40: D1178–D1186.

Gruner P, Miedaner T. 2021. Perennial Rye: Genetics of Perenniality and Limited Fertility. Plants 10: 1210.

Guo K, Liu M, Vella D, Suresh S, Hsia KJ. 2024. Dehydration-induced corrugated folding in Rhapis excelsa plant leaves. Proceedings of the National Academy of Sciences 121: e2320259121.

Haas B. 2023. TransDecoder (Find Coding Regions Within Transcripts).

Hajjar R, Hodgkin T. 2007. The use of wild relatives in crop improvement: a survey of developments over the last 20 years. Euphytica 156: 1–13.

Harlan JR, de Wet JMJ. 1971. Toward a Rational Classification of Cultivated Plants. Taxon 20: 509–517.

Hernandez J, Meints B, Hayes P. 2020. Introgression Breeding in Barley: Perspectives and Case Studies. Frontiers in Plant Science 11.

Hernández-Sánchez IE, Maruri-López I, Martinez-Martinez C, Janis B, Jiménez-Bremont JF, Covarrubias AA, Menze MA, Graether SP, Thalhammer A. 2022. LEAfing through literature: late embryogenesis abundant proteins coming of age—achievements and perspectives. Journal of Experimental Botany 73: 6525–6546.

Holst F, Bolger A, Günther C, Maß J, Triesch S, Kindel F, Kiel N, Saadat N, Ebenhöh O, Usadel B, et al. 2023. Helixer–de novo Prediction of Primary Eukaryotic Gene Models Combining Deep Learning and a Hidden Markov Model. : 2023.02.06.527280.

Hübner S, Höffken M, Oren E, Haseneyer G, Stein N, Graner A, Schmid K, Fridman E. 2009. Strong correlation of wild barley (Hordeum spontaneum) population structure with temperature and precipitation variation. Molecular Ecology 18: 1523–1536.

Hundertmark M, Hincha DK. 2008. LEA (Late Embryogenesis Abundant) proteins and their encoding genes in Arabidopsis thaliana. BMC Genomics 9: 1–22.

Hyndman RJ, Einbeck J, Wand MP. 2023. hdrcde: Highest Density Regions and Conditional Density Estimation.

Jackman SD, Coombe L, Chu J, Warren RL, Vandervalk BP, Yeo S, Xue Z, Mohamadi H, Bohlmann J, Jones SJM, et al. 2018. Tigmint: correcting assembly errors using linked reads from large molecules. BMC Bioinformatics 19: 1–10.

Jakob SS, Blattner FR. 2006. A Chloroplast Genealogy of Hordeum (Poaceae): Long-Term Persisting Haplotypes, Incomplete Lineage Sorting, Regional Extinction, and the Consequences for Phylogenetic Inference. Molecular Biology and Evolution 23: 1602–1612.

Jakob SS, Meister A, Blattner FR. 2004. The Considerable Genome Size Variation of Hordeum Species (Poaceae) Is Linked to Phylogeny, Life Form, Ecology, and Speciation Rates. Molecular Biology and Evolution 21: 860–869.

Jarzyniak KM, Jasiński M. 2014. Membrane transporters and drought resistance – a complex issue. Frontiers in Plant Science 5.

Jayakodi M, Lu Q, Pidon H, Rabanus-Wallace MT, Bayer M, Lux T, Guo Y, Jaegle B, Badea A, Bekele W, et al. 2024. Structural variation in the pangenome of wild and domesticated barley. Nature 636: 654–662.

Jayakodi M, Padmarasu S, Haberer G, Bonthala VS, Gundlach H, Monat C, Lux T, Kamal N, Lang D, Himmelbach A, et al. 2020. The barley pan-genome reveals the hidden legacy of mutation breeding. Nature 588: 284–289.

Jones P, Binns D, Chang H-Y, Fraser M, Li W, McAnulla C, McWilliam H, Maslen J, Mitchell A, Nuka G, et al. 2014. InterProScan 5: genome-scale protein function classification. Bioinformatics 30: 1236–1240.

Kadioglu A, Terzi R, Saruhan N, Saglam A. 2012. Current advances in the investigation of leaf rolling caused by biotic and abiotic stress factors. Plant Science 182: 42–48.

Kasapligil B. 1961. Foliar Xeromorphy of Certain Geophytic Monocotyledons. Madroño 16: 43–70.

Kashyap A, Garg P, Tanwar K, Sharma J, Gupta NC, Ha PTT, Bhattacharya RC, Mason AS, Rao M. 2022. Strategies for utilization of crop wild relatives in plant breeding programs. Theoretical and Applied Genetics 135: 4151–4167.

Kolmogorov M, Yuan J, Lin Y, Pevzner PA. 2019. Assembly of long, error-prone reads using repeat graphs. Nature Biotechnology 37: 540–546.

von Korff M, Wang H, Léon J, Pillen K. 2004. Development of candidate introgression lines using an exotic barley accession (Hordeum vulgare ssp. spontaneum) as donor. Theoretical and Applied Genetics 109: 1736–1745.

Kumar S, Suleski M, Craig JM, Kasprowicz AE, Sanderford M, Li M, Stecher G, Hedges SB. 2022. TimeTree 5: An Expanded Resource for Species Divergence Times. Molecular Biology and Evolution 39: msac174.

Li H. 2018. Minimap2: pairwise alignment for nucleotide sequences. Bioinformatics 34: 3094–3100.

Liakoura V, Stefanou M, Manetas Y, Cholevas C, Karabourniotis G. 1997. Trichome density and its UV-B protective potential are affected by shading and leaf position on the canopy. Environmental and Experimental Botany 38: 223–229.

Liao Y, Smyth GK, Shi W. 2014. featureCounts: an efficient general purpose program for assigning sequence reads to genomic features. Bioinformatics 30: 923–930.

Liller CB, Walla A, Boer MP, Hedley P, Macaulay M, Effgen S, von Korff M, van Esse GW, Koornneef M. 2017. Fine mapping of a major QTL for awn length in barley using a multiparent mapping population. Theoretical and Applied Genetics 130: 269–281.

Lisch D. 2013. How important are transposons for plant evolution? Nature Reviews Genetics 14: 49– 61.

Liu Y, Du H, Li P, Shen Y, Peng H, Liu S, Zhou G-A, Zhang H, Liu Z, Shi M, et al. 2020. Pan-Genome of Wild and Cultivated Soybeans. Cell 182: 162–176.e13.

Liu X, Wang Z, Wang L, Wu R, Phillips J, Deng X. 2009. *LEA* 4 group genes from the resurrection plant *Boea hygrometrica* confer dehydration tolerance in transgenic tobacco. Plant Science 176: 90– 98.

Ma J, Bennetzen JL. 2004. Rapid recent growth and divergence of rice nuclear genomes. Proceedings of the National Academy of Sciences 101: 12404–12410.

Manni M, Berkeley MR, Seppey M, Simão FA, Zdobnov EM. 2021. BUSCO Update: Novel and Streamlined Workflows along with Broader and Deeper Phylogenetic Coverage for Scoring of Eukaryotic, Prokaryotic, and Viral Genomes. Molecular Biology and Evolution 38: 4647–4654.

Marçais G, Kingsford C. 2011. A fast, lock-free approach for efficient parallel counting of occurrences of k-mers. Bioinformatics 27: 764–770.

Marks RA, Van Der Pas L, Schuster J, Gilman IS, VanBuren R. 2024. Convergent evolution of desiccation tolerance in grasses. Nature Plants 10: 1112–1125.

Mascher M, Marone MP, Schreiber M, Stein N. 2024. Are cereal grasses a single genetic system? Nature Plants 10: 719–731.

Mascher M, Wicker T, Jenkins J, Plott C, Lux T, Koh CS, Ens J, Gundlach H, Boston LB, Tulpová Z, et al. 2021. Long-read sequence assembly: a technical evaluation in barley. The Plant Cell 33: 1888–1906.

Matus I, Corey A, Filichkin T, Hayes PM, Vales MI, Kling J, Riera-Lizarazu O, Sato K, Powell W, Waugh R. 2003. Development and characterization of recombinant chromosome substitution lines (RCSLs) using Hordeum vulgare subsp. spontaneum as a source of donor alleles in a Hordeum vulgare subsp. vulgare background. Genome 46: 1010–1023.

Mazel A, Leshem Y, Tiwari BS, Levine A. 2004. Induction of Salt and Osmotic Stress Tolerance by Overexpression of an Intracellular Vesicle Trafficking Protein AtRab7 (AtRabG3e). Plant Physiology 134: 118–128.

Mendes FK, Vanderpool D, Fulton B, Hahn MW. 2021. CAFE 5 models variation in evolutionary rates among gene families. Bioinformatics 36: 5516–5518.

Mendiburu F de. 2023. agricolae: Statistical Procedures for Agricultural Research.

Middleton CP, Stein N, Keller B, Kilian B, Wicker T. 2013. Comparative analysis of genome composition in Triticeae reveals strong variation in transposable element dynamics and nucleotide diversity. The Plant Journal 73: 347–356.

Mikheenko A, Prjibelski A, Saveliev V, Antipov D, Gurevich A. 2018. Versatile genome assembly evaluation with QUAST-LG. Bioinformatics 34: i142–i150.

Mittler R, Finka A, Goloubinoff P. 2012. How do plants feel the heat? Trends in Biochemical Sciences 37: 118–125.

Mittler R, Vanderauwera S, Suzuki N, Miller G, Tognetti VB, Vandepoele K, Gollery M, Shulaev V, Van Breusegem F. 2011. ROS signaling: the new wave? Trends in Plant Science 16: 300–309.

Monat C, Padmarasu S, Lux T, Wicker T, Gundlach H, Himmelbach A, Ens J, Li C, Muehlbauer GJ, Schulman AH, et al. 2019. TRITEX: chromosome-scale sequence assembly of Triticeae genomes with open-source tools. Genome Biology 20: 1–18.

Nagy Z, Németh E, Guóth A, Bona L, Wodala B, Pécsváradi A. 2013. Metabolic indicators of drought stress tolerance in wheat: Glutamine synthetase isoenzymes and Rubisco. Plant Physiology and Biochemistry 67: 48–54.

Navrátilová P, Toegelová H, Tulpová Z, Kuo Y-T, Stein N, Doležel J, Houben A, Šimková H, Mascher M. 2022. Prospects of telomere-to-telomere assembly in barley: Analysis of sequence gaps in the MorexV3 reference genome. Plant Biotechnology Journal 20: 1373–1386.

Nelson DE, Repetti PP, Adams TR, Creelman RA, Wu J, Warner DC, Anstrom DC, Bensen RJ, Castiglioni PP, Donnarummo MG, et al. 2007. Plant nuclear factor Y (NF-Y) B subunits confer drought tolerance and lead to improved corn yields on water-limited acres. Proceedings of the National Academy of Sciences 104: 16450–16455.

Neumann P, Navrátilová A, Koblížková A, Kejnovský E, Hřibová E, Hobza R, Widmer A, Doležel J, Macas J. 2011. Plant centromeric retrotransposons: a structural and cytogenetic perspective. Mobile DNA 2: 4.

Nieves-Cordones M, García-Sánchez F, Pérez-Pérez JG, Colmenero-Flores JM, Rubio F, Rosales MA. 2019. Coping With Water Shortage: An Update on the Role of K+, Cl-, and Water Membrane Transport Mechanisms on Drought Resistance. Frontiers in Plant Science 10.

Nigel Maxted, Laura Rhodes, Isabelle Bradley. 2014. IUCN Red List of Threatened Species: Hordeum erectifolium. IUCN Red List of Threatened Species.

Noctor G, Mhamdi A, Foyer CH. 2014. The Roles of Reactive Oxygen Metabolism in Drought: Not So Cut and Dried. Plant Physiology 164: 1636–1648.

Nueda MJ, Tarazona S, Conesa A. 2014. Next maSigPro: updating maSigPro bioconductor package for RNA-seq time series. Bioinformatics 30: 2598–2602.

Onoda Y, Wright IJ, Evans JR, Hikosaka K, Kitajima K, Niinemets Ü, Poorter H, Tosens T, Westoby M. 2017. Physiological and structural tradeoffs underlying the leaf economics spectrum. New Phytologist 214: 1447–1463.

Ou S, Jiang N. 2018. LTR_retriever: A Highly Accurate and Sensitive Program for Identification of Long Terminal Repeat Retrotransposons. Plant Physiology 176: 1410–1422.

Ou S, Su W, Liao Y, Chougule K, Agda JRA, Hellinga AJ, Lugo CSB, Elliott TA, Ware D, Peterson T, et al. 2019. Benchmarking transposable element annotation methods for creation of a streamlined, comprehensive pipeline. Genome Biology 20: 1–18.

Pankin A, Altmüller J, Becker C, von Korff M. 2018. Targeted resequencing reveals genomic signatures of barley domestication. New Phytologist 218: 1247–1259.

Pardo J, Man Wai C, Chay H, Madden CF, Hilhorst HWM, Farrant JM, VanBuren R. 2020. Intertwined signatures of desiccation and drought tolerance in grasses. Proceedings of the National Academy of Sciences 117: 10079–10088.

Peleg Z, Reguera M, Tumimbang E, Walia H, Blumwald E. 2011. Cytokinin-mediated source/sink modifications improve drought tolerance and increase grain yield in rice under water-stress. Plant Biotechnology Journal 9: 747–758.

Perico C, Tan S, Langdale JA. 2022. Developmental regulation of leaf venation patterns: monocot versus eudicots and the role of auxin. New Phytologist 234: 783–803.

Poorter H, Niinemets Ü, Poorter L, Wright IJ, Villar R. 2009. Causes and consequences of variation in leaf mass per area (LMA): a meta-analysis. New Phytologist 182: 565–588.

Presting GG, Malysheva L, Fuchs J, Schubert I. 1998. A TY3/GYPSY retrotransposon-like sequence localizes to the centromeric regions of cereal chromosomes. The Plant Journal 16: 721– 728.

Prjibelski AD, Mikheenko A, Joglekar A, Smetanin A, Jarroux J, Lapidus AL, Tilgner HU. 2023. Accurate isoform discovery with IsoQuant using long reads. Nature Biotechnology 41: 915– 918.

R Core Team. 2024. R: A Language and Environment for Statistical Computing. Vienna, Austria: R Foundation for Statistical Computing.

Redmann RE. 1985. Adaptation of Grasses to Water Stress-Leaf Rolling and Stomate Distribution. Annals of the Missouri Botanical Garden 72: 833–842.

Renzi JP, Coyne CJ, Berger J, von Wettberg E, Nelson M, Ureta S, Hernández F, Smýkal P, Brus J. 2022. How Could the Use of Crop Wild Relatives in Breeding Increase the Adaptation of Crops to Marginal Environments? Frontiers in Plant Science 13.

Rhie A, Walenz BP, Koren S, Phillippy AM. 2020. Merqury: reference-free quality, completeness, and phasing assessment for genome assemblies. Genome Biology 21: 1–27.

Sakuma Y, Maruyama K, Osakabe Y, Qin F, Seki M, Shinozaki K, Yamaguchi-Shinozaki K. 2006. Functional Analysis of an Arabidopsis Transcription Factor, DREB2A, Involved in Drought-Responsive Gene Expression. The Plant Cell 18: 1292–1309.

Sgroi LC, Lovino MA, Berbery EH, Müller GV. 2021. Characteristics of droughts in Argentina’s core crop region. Hydrology and Earth System Sciences 25: 2475–2490.

Shields LM. 1950. Leaf Xeromorphy as Related to Physiological and Structural Influences. Botanical Review 16: 399–447.

Šimková H, Tulpová Z, Cápal P. 2023. Flow Sorting–Assisted Optical Mapping. In: Heitkam T, Garcia S, eds. Plant Cytogenetics and Cytogenomics: Methods and Protocols. New York, NY: Springer US, 465–483.

Skirycz A, Inzé D. 2010. More from less: plant growth under limited water. Current Opinion in Biotechnology 21: 197–203.

Skirycz A, Vandenbroucke K, Clauw P, Maleux K, De Meyer B, Dhondt S, Pucci A, Gonzalez N, Hoeberichts F, Tognetti VB, et al. 2011. Survival and growth of Arabidopsis plants given limited water are not equal. Nature Biotechnology 29: 212–214.

Stritt C, Wyler M, Gimmi EL, Pippel M, Roulin AC. 2020. Diversity, dynamics and effects of long terminal repeat retrotransposons in the model grass Brachypodium distachyon. New Phytologist 227: 1736–1748.

Sun J, Cui X, Teng S, Kunnong Z, Wang Y, Chen Z, Sun X, Wu J, Ai P, Quick WP, et al. 2020. HD-ZIP IV gene Roc8 regulates the size of bulliform cells and lignin content in rice. Plant Biotechnology Journal 18: 2559–2572.

Sun H, Ding J, Piednoël M, Schneeberger K. 2018. findGSE: estimating genome size variation within human and Arabidopsis using k-mer frequencies. Bioinformatics 34: 550–557.

Sun Y, McManus JF, Clemens SC, Zhang X, Vogel H, Hodell DA, Guo F, Wang T, Liu X, An Z. 2021. Persistent orbital influence on millennial climate variability through the Pleistocene. Nature Geoscience 14: 812–818.

The UniProt Consortium, Bateman A, Martin M-J, Orchard S, Magrane M, Ahmad S, Alpi E, Bowler-Barnett EH, Britto R, Bye-A-Jee H, et al. 2023. UniProt: the Universal Protein Knowledgebase in 2023. Nucleic Acids Research 51: D523–D531.

VanBuren R, Pardo J, Man Wai C, Evans S, Bartels D. 2019. Massive Tandem Proliferation of ELIPs Supports Convergent Evolution of Desiccation Tolerance across Land Plants. Plant Physiology 179: 1040–1049.

Vaser R, Sović I, Nagarajan N, Šikić M. 2017. Fast and accurate de novo genome assembly from long uncorrected reads. Genome Research 27: 737–746.

Vasimuddin Md, Misra S, Li H, Aluru S. 2019. Efficient Architecture-Aware Acceleration of BWA-MEM for Multicore Systems. In: 2019 IEEE International Parallel and Distributed Processing Symposium (IPDPS). 314–324.

Vincent H, Bothmer R von, Knüpffer H, Amri A, Konopka J, Maxted N. 2012. Genetic gap analysis of wild Hordeum taxa. Plant Genetic Resources 10: 242–253.

Vuilleumier BS. 1971. Pleistocene Changes in the Fauna and Flora of South America. Science 173: 771–780.

Waddington SR, Cartwright PM, WALL PC. 1983. A Quantitative Scale of Spike Initial and Pistil Development in Barley and Wheat. Annals of Botany 51: 119–130.

Walther U, Rapke H, Proeseler G, Szigat G. 2000. Hordeum bulbosum-a new source of disease resistance-transfer of resistance to leaf rust and mosaic viruses from H. bulbosum into winter barley. Plant Breeding 119: 215–218.

Wang H, Lu S, Guan X, Jiang Y, Wang B, Hua J, Zou B. 2022. Dehydration-Responsive Element Binding Protein 1C, 1E, and 1G Promote Stress Tolerance to Chilling, Heat, Drought, and Salt in Rice. Frontiers in Plant Science 13.

Wendler N, Mascher M, Himmelbach A, Johnston P, Pickering R, Stein N. 2015. Bulbosum to Go: A Toolbox to Utilize Hordeum vulgare/bulbosum Introgressions for Breeding and Beyond. Molecular Plant 8: 1507–1519.

Wick RR, Judd LM, Gorrie CL, Holt KE. 2017. Completing bacterial genome assemblies with multiplex MinION sequencing. Microbial Genomics 3.

Wicker T, Gundlach H, Spannagl M, Uauy C, Borrill P, Ramírez-González RH, De Oliveira R, Mayer KFX, Paux E, Choulet F. 2018. Impact of transposable elements on genome structure and evolution in bread wheat. Genome Biology 19: 1–18.

Wickham H, Averick M, Bryan J, Chang W, McGowan LD, François R, Grolemund G, Hayes A, Henry L, Hester J, et al. 2019. Welcome to the Tidyverse. Journal of Open Source Software 4: 1686.

Winterfeld G, Tkach N, Röser M. 2025. Reductional dysploidy and genome size diversity in Pooideae, the largest subfamily of grasses (Poaceae). Plant Systematics and Evolution 311: 18.

WMO. 2020. WMO Climatological Normals | World Meteorological Organization.

Wu H-J, Zhang Z, Wang J-Y, Oh D-H, Dassanayake M, Liu B, Huang Q, Sun H-X, Xia R, Wu Y, et al. 2012. Insights into salt tolerance from the genome of Thellungiella salsuginea. Proceedings of the National Academy of Sciences 109: 12219–12224.

Xu S, Hu E, Cai Y, Xie Z, Luo X, Zhan L, Tang W, Wang Q, Liu B, Wang R, et al. 2024. Using clusterProfiler to characterize multiomics data. Nature Protocols 19: 3292–3320.

Xue D, Zhang X, Lu X, Chen G, Chen Z-H. 2017. Molecular and Evolutionary Mechanisms of Cuticular Wax for Plant Drought Tolerance. Frontiers in Plant Science 8.

Ye Z, Sawada M, Iwasa M, Moriyama R, Dey D, Furutani M, Kitao M, Hara T, Tanaka A, Kishimoto J, et al. 2024. Revisiting the early light-induced protein hypothesis in the sustained thermal dissipation mechanism in yew leaves. Journal of Experimental Botany: erae412.

Zhang H, Dawe RK. 2012. Total centromere size and genome size are strongly correlated in ten grass species. Chromosome Research 20: 403–412.

Zhang S, Huang G, Zhang Y, Lv X, Wan K, Liang J, Feng Y, Dao J, Wu S, Zhang L, et al. 2023. Sustained productivity and agronomic potential of perennial rice. Nature Sustainability 6: 28–38.

Zhang R-G, Li G-Y, Wang X-L, Dainat J, Wang Z-X, Ou S, Ma Y. 2022. TEsorter: An accurate and fast method to classify LTR-retrotransposons in plant genomes. Horticulture Research 9: uhac017.

Zhu Z, Wang J, Li C, Li L, Mao X, Hu G, Wang J, Chang J, Jing R. 2022. A transcription factor TaMYB5 modulates leaf rolling in wheat. Frontiers in Plant Science 13.

